# hGRAD – a versatile ʹone-fits-allʹ system for the acute depletion of RNA binding proteins in nuclear condensates

**DOI:** 10.1101/2023.08.04.551933

**Authors:** Benjamin Arnold, Ricarda J. Riegger, Ellen Kazumi Okuda, Irena Slišcović, Mario Keller, François McNicoll, Kathi Zarnack, Michaela Müller-McNicoll

## Abstract

Nuclear RNA binding proteins (RBPs) are difficult to study because they often belong to large protein families and form extensive networks of auto- and cross-regulation. They are highly abundant and often localize to condensates with a slow turnover, requiring long depletion times or knockouts that cannot distinguish between direct and indirect or compensatory effects.

Here, we developed a system that is optimized for the rapid degradation of nuclear RBPs, called hGRAD. It comes as a ʹone-fits-allʹ plasmid, and integration into any cell line that expresses endogenously GFP-tagged proteins allows an inducible, rapid and complete knockdown. We show that the nuclear RBPs SRSF3, SRSF5, SRRM2 and NONO are completely cleared from nuclear speckles and paraspeckles within two hours. hGRAD works in various cell types, is more efficient than other methods and does not require the expression of exogenous ubiquitin ligases. Combining SRSF5 hGRAD degradation with Nascent-seq uncovered highly dynamic transient transcript changes, compensatory mechanisms and that SRSF5 promotes transcript stability.

## Introduction

RNA binding proteins (RBPs) often belong to larger protein families and form dense regulatory networks of auto- and cross-regulation, wherein they complement or antagonize each otherʹs functions. These networks ensure the robustness of gene expression starting from transcription in the nucleus to translation and mRNA decay in the cytoplasm. This renders it very challenging to disentangle the specific functions of individual RBPs and their relationship to human diseases (Gebauer et al., 2021; Müller- McNicoll et al., 2019). Most RBPs are also highly abundant and stable proteins, and many of them localize to nuclear condensates with a slow turnover, which makes it difficult to deplete them using RNA interference. On the other hand, complete knockouts (KOs) or very long depletion times trigger compensatory effects and do not allow to distinguish between direct and indirect effects.

One example for such a large family of nuclear RBPs are the serine/arginine-rich splicing factors (SRSF1-SRSF12, SR proteins) that are present in all metazoans (Sliskovic et al., 2022). SR proteins are essential factors for constitutive and alternative pre-mRNA splicing (AS), and they also influence many other steps in mRNA metabolism including transcription, alternative polyadenylation, mRNA packaging, export, translation and cytoplasmic decay (Jeong, 2017; Wegener and Müller-McNicoll, 2019; Änkö, 2014). SR proteins also participate in the processing of non-coding RNAs and the formation and dynamics of nuclear compartments (de Oliveira Freitas Machado et al., 2023; Königs et al., 2020; Wagner and Frye, 2021). In contrast to their canonical functions in splicing, the participation of individual SR protein family members in specific AS events or other disease-related, non-canonical functions remains poorly understood. This is in part because SR- and SR-like proteins cooperate and compete for binding sites, compensate for each other, and the removal of one SR protein can affect the levels of several other family members (Meinke et al., 2020; Änkö et al., 2012). Dysregulated levels of individual SR protein family members are associated with various cancers and other diseases including neurological disorders, liver disease as well as coronary and cardiac diseases (More and Kumar, 2020; Ratnadiwakara et al., 2018; Zheng et al., 2020). Because of their overlapping functions and compensatory mechanisms, specific inducible protein degradation systems with short depletion times are needed to disentangle the functions of individual SR proteins and other RBPs.

Proteins are usually targeted for degradation by the addition of polyubiquitin chains to lysine residues which are recognized by the 26S proteasome (Komander and Rape, 2012; Yu and Matouschek, 2017). In a highly regulated multi-enzymatic reaction, ubiquitin (Ub) is first activated by E1 enzymes under ATP consumption, subsequently transferred to E2-Ub-conjugation enzymes and finally transferred to target proteins by E3-Ub**-**ligases. More than 600 different E3-Ub-ligases are encoded in the human genome, enabling target specificity (Nakayama and Nakayama, 2006). The RING-finger family contains the SKP1-Cullin-F-box protein (SCF) complex, which is the most prominent E3- Ub-ligase in mammals (Frescas and Pagano, 2008; Schulman et al., 2000). Its subunit RBX1 facilitates E2-Ub recruitment and the ubiquitylation reaction, CUL1 functions as a bridge and SKP1 mediates binding to a modular F-box protein (FBP). More than 70 different FBPs are encoded in the human genome, providing additional target specificity (Frescas and Pagano, 2008; Schulman et al., 2000). FBPs have two major domains: the F-box domain, which is essential for the interaction with SKP1, and a variable C-terminal domain, which is essential for target recognition (Kipreos and Pagano, 2000).

Most targeted protein degradation approaches manipulate the SCF complex by introducing an engineered FBP with specificity for a protein-of-interest (POI). Examples are the deGradFP system (Caussinus et al., 2011) and the auxin-inducible degron (AID) systems (Li et al., 2019; Nishimura et al., 2009). The deGradFP system was developed to degrade GFP-tagged proteins in *Drosophila melanogaster.* Here, the substrate recognition of the F-box protein Slmb was altered by fusing its minimal F-box to a synthetic anti-GFP nanobody (VHH-GFP4) (Rothbauer et al., 2008) replacing the C-terminal WD40 repeat domain (Caussinus et al., 2011). Nanobodies are small proteins (10-15 kDa) that resemble the variable fragment (VHH) of homodimeric heavy chain antibodies, derived from camelids (Harmsen and De Haard, 2007). They are highly stable, can be easily expressed in mammalian cells and bind strongly to their specific antigens (Beghein and Gettemans, 2017). Upon inducible expression of the F-box-nanobody fusion protein, the GFP-tagged POI is bound, endogenous SKP1, CUL1 and RBX1 are recruited, and ubiquitylation and degradation are initiated. The deGradFP system degrades GFP-tagged proteins in *Drosophila* with a half-life of 2 hours and is also functional for several GFP derivates like Venus, YFP and EYFP (Caussinus and Affolter, 2016; Caussinus et al., 2011). It was successfully adapted to other model organisms, such as *Danio rerio* and *Trypanosoma brucei* (Ishii and Akiyoshi, 2022; Yamaguchi et al., 2019), but so far failed to degrade nuclear proteins in mammalian cells (Daniel et al., 2018; Ludwicki et al., 2019; Shin et al., 2015).

In the AID systems, the plant hormone auxin (indole-3-acetic acid, IAA or AUX) induces target protein degradation via the SCFs osTIR1 from *Oriza sativa* (Nishimura et al., 2009) or aaAFB2 from *Arabidopsis thaliana* (Li et al., 2019). Upon AUX binding, osTIR1 undergoes a structural change allowing stable interactions with AID sequences in target proteins, which are subsequently ubiquitylated and degraded (Holland et al., 2012; Nishimura et al., 2009; Trost et al., 2016). AID sequences can be fused directly to the target proteins or to nanobodies, which can be inducibly expressed and bind to the target protein. Upon addition of AUX, degradation occurs within 20-60 minutes, and is stopped upon AUX withdrawal (Daniel et al., 2018; Holland et al., 2012; Li et al., 2019; Nishimura et al., 2009). Despite these advantages, AID systems require the constitutive expression of large foreign FBPs, which is often toxic for cells, and they are leaky showing basal degradation without auxin induction.

An alternative degradation system that functions independently of the SCF complex is called Trim-Away and is based on the E3-Ub-ligase tripartite motif-containing protein 21 (TRIM21), which confers target specificity and E3-Ub-ligase activity (Clift et al., 2017). TRIM21 recognizes and binds the Fc region of antigen-bound antibodies (IgG) and mediates their ubiquitylation and degradation (James et al., 2007). This system enables a rapid and efficient degradation of cytoplasmic proteins, with half-lives of 10-30 minutes. However, target-specific antibodies need to be delivered into cells by electroporation which, due to their large size, cannot diffuse through the nuclear pore and thus cannot be used to deplete nuclear proteins (Clift et al., 2017).

Here, we present hGRAD, short for <u>h</u>uman protein de<u>grad</u>ation system, which is optimized to induce the rapid and efficient degradation of GFP-tagged nuclear RBPs from various subnuclear compartments. hGRAD allowed us to identify a novel function of SRSF5 in promoting transcript stability and to define direct target genes and compensatory mechanisms.

## Results

### Development of the hGRAD system

To study the functions of nuclear RBPs, we aimed to develop a rapid degradation system that is easy to implement, requires minimal engineering of host cells and efficiently degrades RBPs that localize to nuclear condensates. Ideally, it should function in different species, cell types and subcellular compartments, and allow to follow the kinetics of degradation in real time. We decided to target GFP-tagged proteins, since many groups use endogenous GFP tagging by CRISPR/Cas9 to study the functions of condensate RBPs and many tagged cell lines are commercially available.

We first designed a ʹone-fits-allʹ plasmid expression system, which allows the inducible expression of two genes in a doxycycline (DOX)-dependent manner (**Figure 1A**). This vector, named pTRE-BI, expresses a Tet-regulator (Tet-on 3G) from the *EF-1α* promoter which drives robust constitutive expression independently of mammalian cell type and species. It also contains a bi-directional tet-inducible promoter (TRE3G BI) with two flanking multiple cloning sites (MCS) and a puromycin resistance gene (PuroR) for genomic integration and selection (**Figure 1A**).

**Figure 1.**
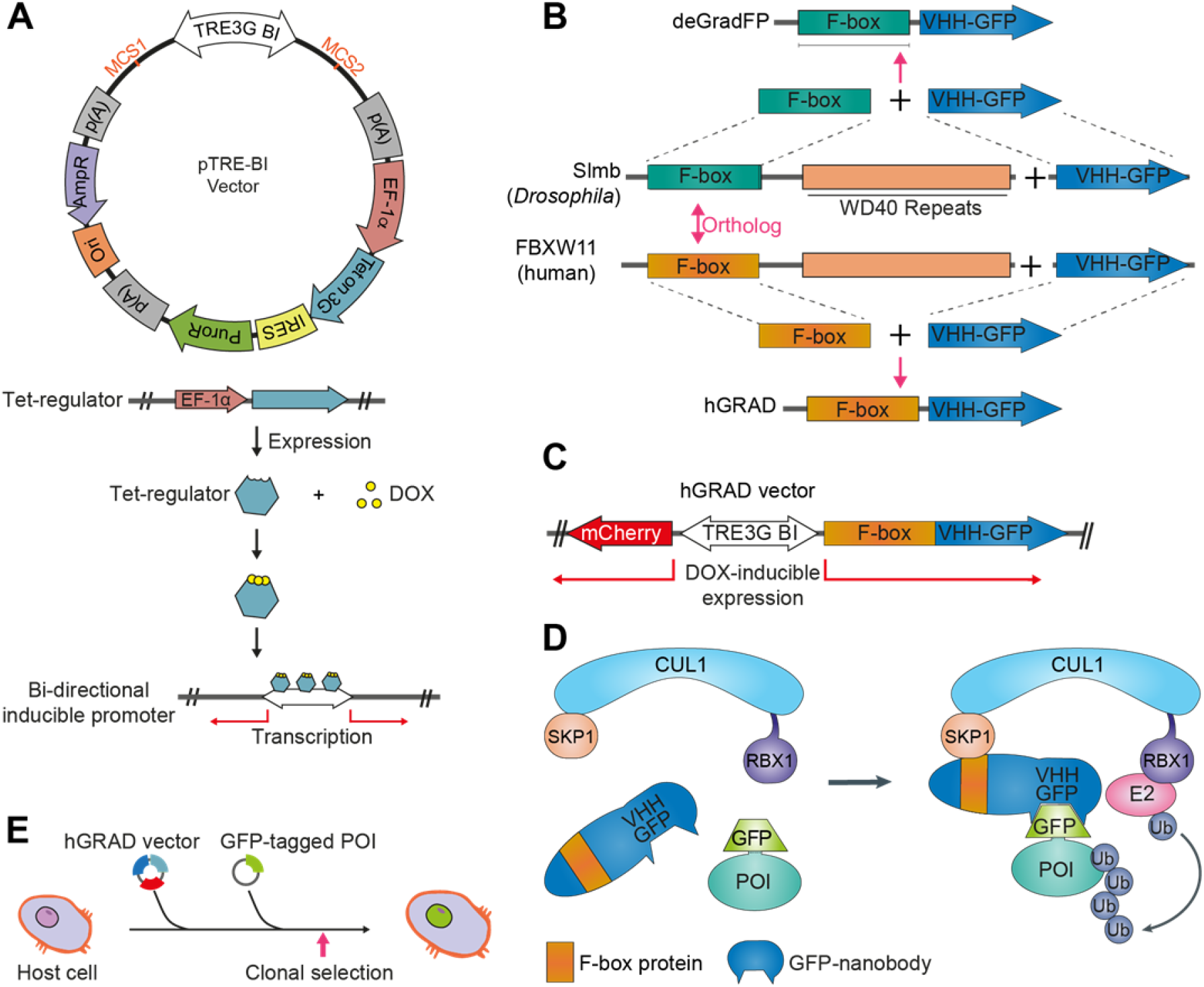
Development of the hGRAD system. **A)** Scheme of the pTRE-BI ʹone-fits-allʹ backbone vector. MCS1 and MCS2, multiple cloning sites; p(A), polyadenylation sites; *EF-1α,* promoter; Tet on 3G, gene encoding Tet-regulator; IRES, internal ribosomal entry site; PuroR, puromycin resistance gene; Ori, origin of replication; AmpR, ampicillin resistance gene. **B)** The hGRAD system is based on the deGradFP system (Caussinus et al., 2011) and was optimized for mammalian protein degradation by exchanging the *Drosophila* Slmb F-box domain with the human FBXW11 F-box domain. **C)** Scheme of the pTRE-BI hGRAD vector. **D)** Overview of the hGRAD system harnessing the cellular SCF complex for ubiquitylation and degradation of the GFP-tagged protein-of-interest (POI). Model modified from (Yamaguchi et al., 2019). **E)** Random genomic integration of the pTRE-BI-hGRAD plasmid allows the inducible degradation of any GFP- tagged protein in any mammalian host cell type.

Next, we modified the deGradFP system (Caussinus et al., 2011), since previous studies indicated that deGradFP is inefficient in degrading nuclear proteins in mammalian cells (Daniel et al., 2018). In humans, more than 70 different FBPs have been reported that mediate target selection and specificity (Frescas and Pagano, 2008; Jin et al., 2004; Zheng et al., 2002). We identified the human protein FBXW11 as the closest ortholog to *Drosophila* Slmb (**Figure 1B**) and fused its minimal F-box domain to an anti- GFP nanobody (VHH-GFP4). The F-box-nanobody fusion was inserted on one side of the bi-directional TREG3 BI promoter and mCherry as induction control on the other side. The Tet-regulator activates the transcription of both genes upon DOX addition. We named our ʹone-fits-allʹ plasmid system hGRAD for <u>h</u>uman protein de<u>grad</u>ation system (**Figure 1C**). Random genomic integration of this vector should allow the inducible degradation of any GFP-tagged protein through binding of the anti-GFP nanobody and recruitment of the cellular SCF complex, which mediates ubiquitylation and proteasomal degradation (**Figure 1D, 1E**).

### hGRAD degrades nuclear RBPs more efficiently than other degradation systems

To be able to compare the performance of hGRAD in degrading mammalian GFP-tagged proteins with other rapid degradation systems, we modified the target specificity of osTIR1, aaAFB2 (AID systems) and TRIM21 (Trim-Away system) through anti-GFP nanobodies and cloned all necessary protein-coding sequences in the pTRE-BI vector (**Figure 2A**): genes encoding *osTIR1* and *aaAFB2* were fused to mCherry and to a weak nuclear localization sequence (NLS) at their C-terminus to ensure optimal degradation of cytoplasmic and nuclear proteins. The gene fusions were then inserted on one side of the bidirectional promoter. On the other side, a sequence encoding a GFP nanobody fused to the minimized AID (mAID) sequence was inserted for the osTIR1 system (Daniel et al., 2018), while for the aaAFB2 system, an optimized AID sequence (miniIAA7) was used (Li et al., 2019). For the Trim-Away system, the *TRIM21* gene was also fused to mCherry and a weak NLS, and the anti-GFP nanobody was fused to the Fc region of human IgG1 (Clift et al., 2017) (hIgG1-Fc2; **Figure 2A**).

**Figure 2.**
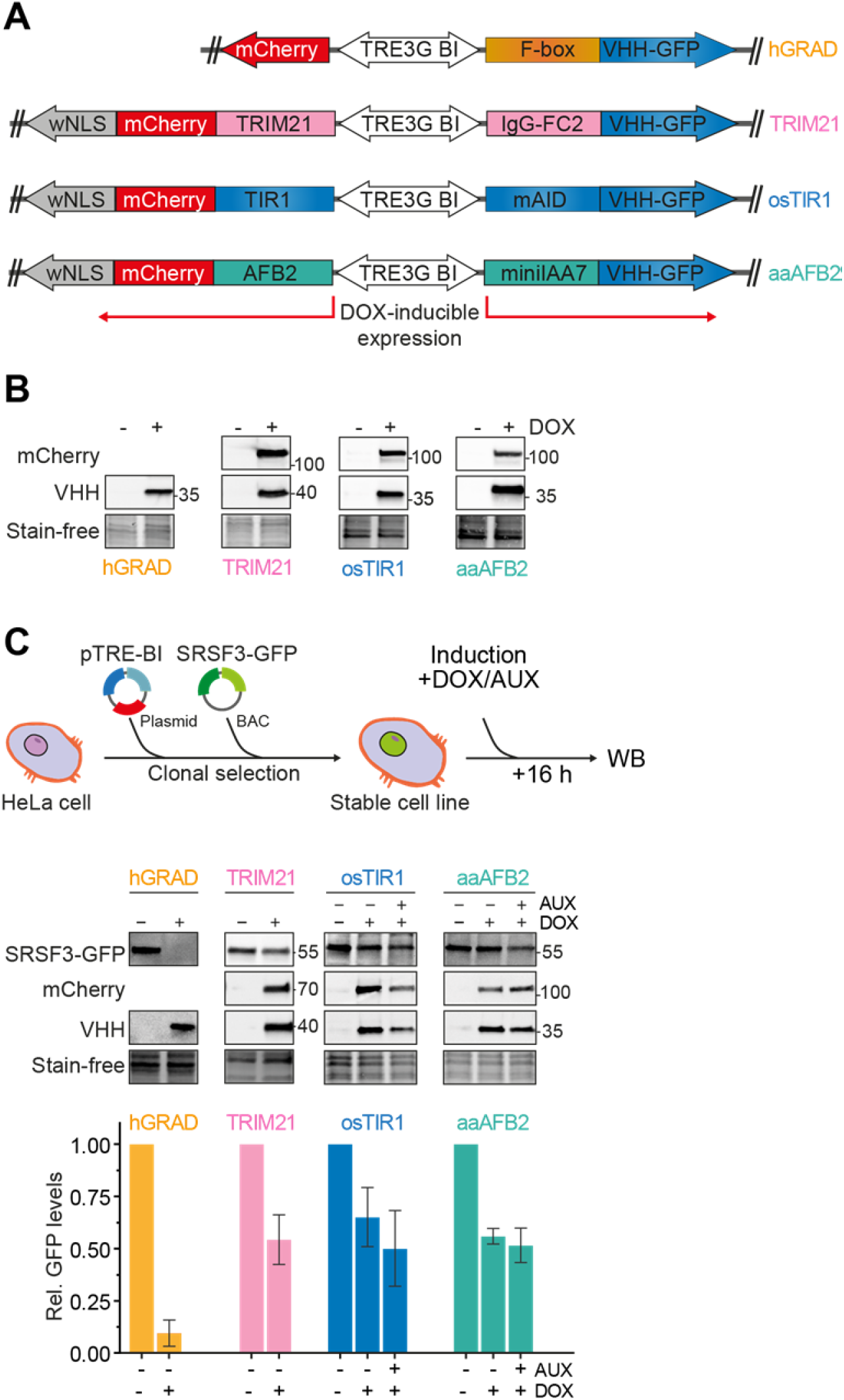
hGRAD degrades the nuclear RBP SRSF3 more efficiently than other degradation systems in HeLa cells. **A)** Schemes of the pTRE-BI ʹone-fits-allʹ vectors for expression of the TRIM21, hGRAD, osTIR1 and aaAFB2 systems. **B)** Induction in HeLa cells (16 h, 1 μg/mL DOX, 0.5 mM AUX). TRIM21-, osTIR1- and aaAFB2-mCherry fusion proteins were detected by an anti-mCherry antibody. The different nanobody fusions (IgG-Fc2-, FBXW11-, mAID- and miniIAA7-VHH-GFP) were detected by an anti-camelid VHH antibody. Stain-free gels were used to detect total protein and control for equal loading. **C)** Experimental scheme and comparison of the degradation efficiencies of TRIM21, hGRAD, osTIR1 or aaAFB2 systems after 16 h DOX and AUX induction. SRSF3-GFP, mCherry and the hGRAD nanobody fusion were detected with anti-GFP, anti-mCherry and anti-camelid antibodies, respectively. Shown are a representative Western blot and the quantification of n=3 independent replicates. Error bars show standard deviation of the mean (SD).

We generated four master cell lines by integrating the different pTRE-BI constructs into the genome of wild-type (WT) HeLa cells and selecting single clones. All pTRE-BI-based induction systems showed a strong DOX-induced expression of the desired proteins and no expression in the uninduced state (**Figure 2B**). Despite their large molecular weights (>100 kDa), all mCherry fusion proteins localized to both the cytoplasm and the nucleus (**Figure S1A**), demonstrating that the weak NLS is functional. Thus, targeted protein degradation should occur in both subcellular compartments (Franic et al., 2021).

We next performed proliferation assays to evaluate potential adverse effects of the DOX treatment. The four master cell lines were grown in the presence or absence of DOX (1 μg/mL; 72 h) and doubling times (t_d_) were calculated. The osTIR1, aaAFB2, TRIM21 and hGRAD cell lines showed similar growth rates compared to WT HeLa cells in the absence of DOX (t_d_=17.23 h), suggesting that the integrated pTRE-BI plasmid itself does not negatively affect cell growth (**Figure S1B**). However, upon DOX addition, the osTIR1 (t_d_=19.3 h), aaAFB2 (t_d_=20.84 h) and TRIM21 (t_d_=18.41 h) cell lines showed reduced growth rates, indicating potential cytotoxic effects from the enzymatic activity or expression of the heterologous proteins. In contrast, the hGRAD cells showed similar doubling times (t_d_=17.24 h) to WT cells, indicating that expression of the hGRAD components does not affect cell growth (**Figure S1B**).

To compare the efficiency in targeting and degrading GFP-tagged SR proteins, we inserted a bacterial artificial chromosome (BAC) harboring the *SRSF3* gene fused to a GFP-tag into the genome of all four master cell lines (**Figure S1C**). The BAC gene harbors the complete *SRSF3* gene, including the endogenous *SRSF3* promoter, untranslated regions (UTRs), exons and introns so that *SRSF3-GFP* is expressed, processed and regulated similarly to the endogenous gene (Müller-McNicoll et al., 2016). All four cell lines showed similar SRSF3-GFP levels and the expected localization to the nucleoplasm and nuclear speckles (**Figure S1D**).

After DOX treatment (16 h), all proteins required for rapid degradation were strongly expressed (**Figure 2C**). The hGRAD system showed an almost complete degradation of SRSF3-GFP (**Figure 2C**). In contrast, the TRIM21, osTIR1 and aaAFB2 systems reduced SRSF3-GFP levels by only about 50% in the presence of AUX (16 h). Moreover, in the osTIR1 and aaAFB2 systems, SRSF3-GFP levels were already reduced in the presence of DOX without AUX, indicating that the AID system is leaky, in line with previous findings (Verma et al., 2020).

Our data demonstrate that the ʹone-fits-allʹ pTRE-BI vector backbone allows a strong and tightly controlled inducible expression of the factors required for degradation of both nuclear and cytoplasmic proteins. Compared to the osTIR1, aaAFB2 and TRIM21 systems, hGRAD showed the most efficient depletion of a nuclear RBP and mildest cytotoxic effects.

### hGRAD efficiently degrades RBPs that localize to nuclear condensates

We next evaluated the capacity of hGRAD to degrade GFP-tagged proteins in different subcellular compartments. For this, we stably integrated BACs expressing eukaryotic translation initiation factor 4E (EIF4E-GFP) and serine/arginine-rich protein-specific kinase 1 (SRPK1-GFP) as cytoplasmic marker proteins, SRSF5-GFP as marker for nuclear speckles and Non-POU domain-containing octamer-binding protein (NONO-GFP) as a marker for paraspeckles into the HeLa hGRAD cell line (**Figure 3A**). The correct subcellular localization of the GFP-tagged proteins was confirmed by confocal microscopy (**Figures 3D, S2A**). Western blot experiments revealed that the hGRAD- mediated degradation worked most efficiently for proteins localizing to nuclear speckles (SRSF3 and SRSF5) or paraspeckles (NONO; > 90%), while cytoplasmic proteins were degraded slightly less efficiently (EIF4E and SRPK1; **Figure 3B**). This was confirmed by confocal microscopy (**Figure S2A**).

**Figure 3.**
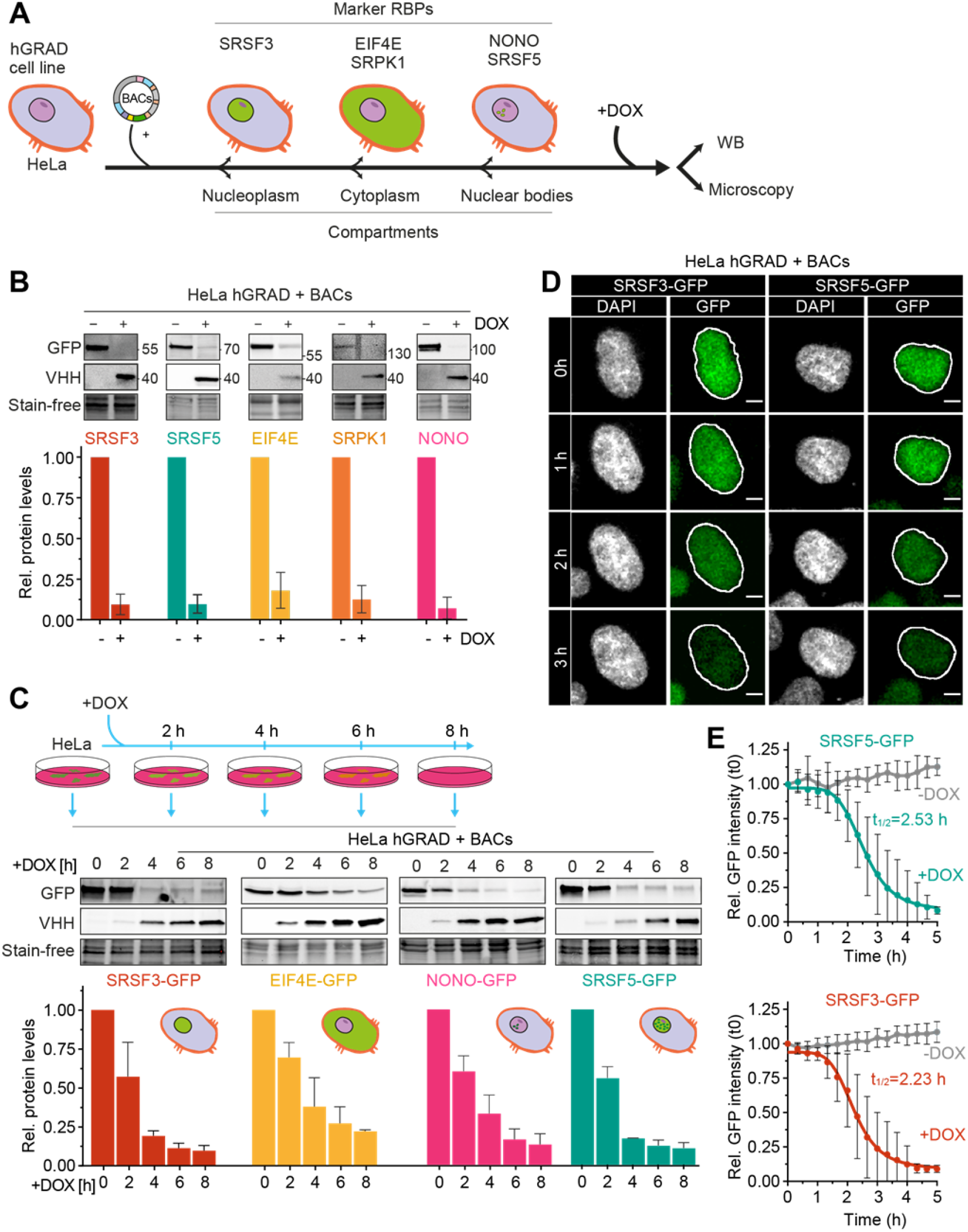
hGRAD works efficiently for RBPs that localize to nuclear condensates. **A)** Experimental scheme to test the hGRAD degradation for human proteins localizing to different subcellular compartments. **B)** Comparison of the degradation efficiency of GFP-tagged RBPs in HeLa cells 16 h after DOX induction (1 μg/mL). **C)** Degradation time-courses show that hGRAD works most efficiently for nuclear RBPs. **B+C)** Shown are the quantifications of n=3 independent Western blot experiments and one representative blot relative to the -DOX or 0 h time point. GFP-tagged proteins and the hGRAD nanobody fusion were detected with anti-GFP and anti-camelid antibody, respectively. **D)** Live-cell imaging to follow the hGRAD degradation kinetics. **E)** Quantification of half-lives (t_1/2_) of SRSF3- and SRSF5-GFP in HeLa cells upon DOX induction. Scale bars = 5 μm. Error bars, SD.

To separate direct from indirect effects, a fast depletion of a protein is most desirable. To evaluate the kinetics of hGRAD-mediated degradation, we performed a time-course for SRSF3, SRSF5, NONO and EIF4E (0, 2, 4, 6 and 8 h DOX; **Figure 3C**). Proteins were already reduced after 2 h post-induction, concomitant with expression of the anti-GFP nanobody. The depletion efficiency differed among the GFP-tagged proteins. Nuclear SRSF3-, SRSF5- and NONO-GFP were degraded rapidly, with half-lives of 2.0 h and 2.5 h and almost 90% depletion after 6 and 8 h respectively. Cytoplasmic EIF4E-GFP was degraded slightly slower, with a half-life of 3.3 h and 75% depletion after 8 h (**Figure 3C, S2B**).

To monitor degradation in individual cells over time, we combined rapid degradation with confocal live-cell imaging. HeLa hGRAD cell lines expressing SRSF3- GFP or SRSF5-GFP were imaged in the presence or absence of DOX for 12 hours. Relative GFP intensity was quantified in individual cells and normalized to the first uninduced time point (t0). Similar to the Western blot quantifications, SRSF3-GFP revealed a half-life of 2.2 h and SRSF5-GFP of 2.5 h (**Figure 3D**). Both proteins reached their minimum levels (below 10% compared to t0) after 4 and 4.5 h of induction, respectively. This indicates that hGRAD degradation of nuclear proteins in individual cells has similar kinetics to bulk cell measurements. Thus, hGRAD quickly and efficiently clears proteins from nuclear condensates within three hours.

### hGRAD works efficiently in different mammalian cell types and species

We next evaluated the capacity of hGRAD to degrade GFP-tagged proteins in pluripotent mouse P19 cells. For this, we generated a P19 hGRAD master cell line and integrated BACs expressing SRSF3-, EIF4E-, SRPK1- and NONO-GFP into the genome (**Figure S2B**). Similar to HeLa cells, the anti-GFP nanobody fusion was well expressed upon DOX induction (1 μg/mL)nand undetectable in the absence of DOX in P19 cells, indicating a strong inducibility and no leaky expression (**Figure 4A**). Proliferation assays revealed that neither integration of the pTRE-BI-hGRAD vector itself (uninduced) nor DOX induction showed any significant changes in growth rates compared to WT P19 cells with the characteristic doubling time of 12 h (**Figure 2B**).

**Figure 4.**
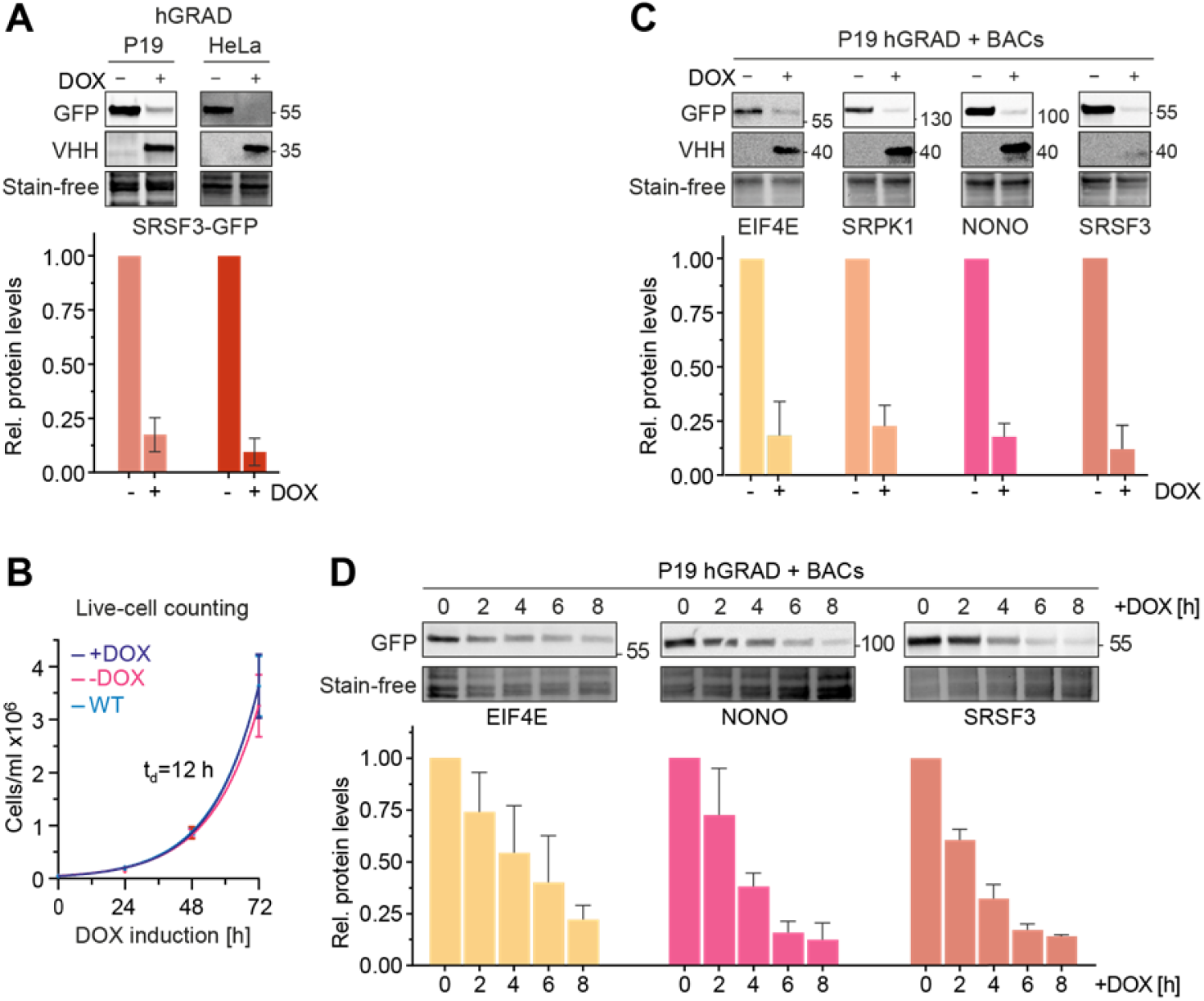
hGRAD works efficiently in different mammalian cell types and species. **A)** Comparison of the degradation efficiency of GFP-tagged SRSF3 in HeLa and P19 cells 16 h after induction with DOX (1 μg/mL). **B)** P19 hGRAD proliferation assay. Growth curves and doubling times (t_d_) were evaluated by exponential growth equation fit (Y=Y_0_*exp(k*X)). P19 WT (light blue), P19 hGRAD -DOX (pink), P19 hGRAD +DOX (dark blue). Shown are mean and SD (error bars) from n=3 independent experiments. **C)** Comparison of the degradation efficiency of GFP-tagged proteins in mouse P19 cells 16 h after DOX induction. **D)** Degradation time-courses show that hGRAD works efficiently in mouse P19 cells. **A-D)** Shown are one representative Western blot and the quantifications of n=3 independent experiments (relative to the -DOX or 0 h time point). GFP-tagged proteins and the hGRAD nanobody fusion were detected with anti-GFP and anti-camelid antibody, respectively. Error bars = SD.

Endpoint degradation assays and GFP fluorescence microscopy confirmed the efficient depletion of all tested proteins in P19 cells (**Figure 4C, S2A**). SRSF3- and NONO-GFP showed a half-life of 2.9 h post-induction and EIF4E-GFP of 4.3 h (**Figure 4D, S2B**). Thus, our data demonstrate that the hGRAD system allows efficient degradation of GFP-tagged proteins in both human HeLa and mouse P19 cells.

### hGRAD works as a rapid knockdown tool for endogenously GFP-tagged nuclear RBPs

We next tested the capacity of hGRAD in degrading endogenously GFP-tagged proteins (endo-GFP) as a rapid knockdown tool. For this, we inserted a GFP-tag at the C-termini of *SRSF3*, *SRSF5* and the nuclear speckle marker *SRRM2* in HeLa and P19 hGRAD master cell lines using CRISPR/Cas9 (**Figure 5A**). Successful GFP-tagging of single clones was validated by PCR, Sanger sequencing and Western blot (**Figure S3A- D**). Confocal microscopy revealed the correct nuclear localization of SRSF3-endo-GFP and SRSF5-endo-GFP, with their typical distribution in the nucleoplasm and enrichment within nuclear speckles, as well as of SRRM2-endo-GFP, which exclusively localized to nuclear speckles (**Figure S3E**). We concluded that all endogenously tagged proteins were correctly folded and fully functional.

**Figure 5.**
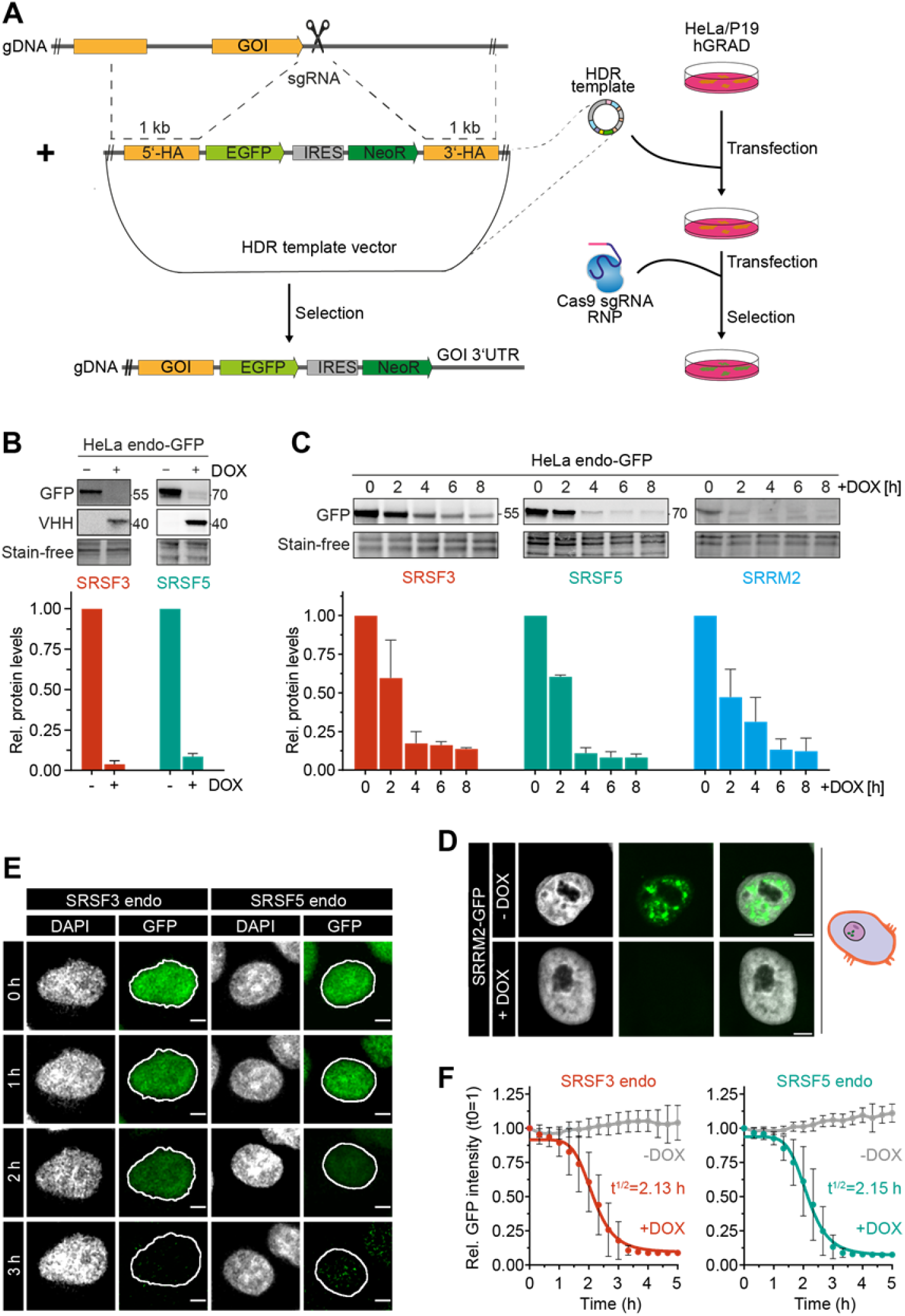
hGRAD efficiently degrades endogenously GFP-tagged nuclear RBPs in HeLa cells allowing their rapid knockdown. **A)** Experimental scheme for the endogenous (endo) GFP-tagging using CRISPR/Cas9 in HeLa and P19 hGRAD cells. gDNA - genomic DNA; sgRNA - single guide RNA; GOI - gene of interest; HDR - homology-directed repair; NeoR - Neomycin resistance cassette; 5ʹ and 3ʹHA - homology arms. **B)** Comparison of the degradation efficiency of SRSF3- and SRSF5-endo-GFP in HeLa cells 16 h after induction by DOX. **C)** Degradation time-courses show that hGRAD works efficiently for SRSF3-, SRSF5- and SRRM2-endo-GFP. **B+C)** Shown are a representative Western blot and the quantification of n=3 independent experiments. Error bars = SD. GFP-tagged proteins and hGRAD nanobody fusion proteins were detected with anti-GFP and anti-camelid antibody, respectively. **D)** Degradation of SRRM2-GFP from nuclear speckles. Scale bars = 5 μm. **E)** Live-cell imaging reveal hGRAD degradation kinetics. **F)** Quantification of the half-lives (t_1/2_) of SRSF3- and SRSF5-endo-GFP in HeLa cells.

Endpoint degradation assays revealed that SRSF3-endo-GFP and SRSF5-endo- GFP were efficiently depleted in HeLa cells (by more than 90%; **Figure 5B**), similar to the system where they were expressed from BACs (see above). Time-course experiments indicated a half-life of 2.2 h for SRSF3-endo-GFP (**Figure 5C**), reaching its minimum after 4 h, similar to SRSF3-GFP from the BAC (2.0 h, **Figure 3C**). SRRM2-endo-GFP showed a half-life of 2.2 h and reached its minimum after 6 h, demonstrating that hGRAD is capable of degrading nuclear proteins that make the core of nuclear speckles (**Figure 5D**). SRSF5-endo-GFP showed a half-life of 2.1 h after DOX induction and reached its minimum levels after 4 h (**Figure 5C**). For comparison, the BAC-expressed SRSF5-GFP showed a half-life of 2.5 h, suggesting that SRSF5-endo-GFP is degraded slightly faster (**Figure 3C, 5C**).

Degradation of SRSF5-endo-GFP was blocked when cells were treated with the proteasome inhibitor MG132 (**Figure S3F**), confirming that degradation occurs via the ubiquitin-proteasome system. In P19 cells, degradation kinetics of SRSF3-endo-GFP and SRSF5-endo-GFP were similar to those in HeLa cells (**Figure S4A, S4B**); both proteins were depleted by 80-90% after 6 h. Live-cell imaging confirmed that SRSF3- and SRSF5- endo-GFP were depleted slightly faster than their counterparts expressed from BACs, with half-lives of 2.13 h and 2.15 h, respectively (**Figure 5E, 5F**).

Interestingly, hGRAD depletion of SRSF5-endo-GFP worked with similar efficiency independently of the glucose concentration (**Figure S4D**), although a previous study proposed that SRSF5 is much less stable under low glucose conditions (Chen et al., 2018). We also did not observe any differences in SRSF5-endo-GFP protein half-lives in high and low glucose conditions (**Figure S4C**). Altogether, our data show that hGRAD efficiently degrades endogenously GFP-tagged SRSF3, SRSF5 and SRRM2, the former in HeLa and P19 cells. This underlines that the simple one-step integration of the hGRAD plasmid into any cell line that expresses endogenously GFP-tagged proteins allows their rapid and inducible knockdown.

### Combining hGRAD with Nascent-seq reveals dynamic changes in transcript levels

Conventional knockdown of SRSF5 using siRNAs required 72 h to obtain a 75% reduction at the protein level (**Figure S5A**). These long depletion times render it difficult to distinguish direct from indirect targets or identify new functions of SRSF5 due to compensatory mechanisms. To improve our understanding of this understudied splicing factor, we combined rapid depletion with nascent RNA sequencing (Nascent-seq) (Radle et al., 2013; Schwalb et al., 2016). Nascent-seq captures only changes that occur in newly transcribed RNAs and ignores the large fraction of pre-existing nuclear and cytoplasmic RNAs that would mask these changes. Especially at early time points, only a small number of differentially expressed or alternatively spliced targets are expected.

Homozygously tagged HeLa SRSF5-endo-GFP cells (**Figure S3D**) were induced with DOX and, at different time points (2, 8, and 16 h; termed T2, T8, and T16, respectively), were treated for one hour with 400 μM 4-thio-uridine (4sU), which is incorporated into newly transcribed RNAs (**Figure 6A**). Uninduced cells were used as controls, and SRSF5 depletion was controlled by Western blot (**Figure S5B**). After total RNA extraction, nascent transcripts with incorporated 4sU were biotinylated *in vitro*, purified using streptavidin-coated beads and converted into a strand-specific cDNA library (**Figure 6A, S5C**). 40% of uniquely mapped reads corresponded to intronic regions, in line with an enrichment of nascent, incompletely spliced pre-mRNAs. Surprisingly, differential expression analysis using DESeq2 (Love et al., 2014) uncovered very rapid and dynamic changes in gene expression after SRSF5 depletion (**Figure 6B, 6C; Table S1**). At T2, already 436 genes were differentially expressed (**Figure 6B**). Of those, 127 were exclusively regulated at T2, whereas 178 differentially expressed genes (DEGs) changed over the entire time-course. Interestingly, at T2, 87% of all DEGs were down-regulated (**Figure S5D**). Among them were 106 (28%) long non-coding RNAs (lncRNAs; **Figure 6C**). 50% of the down-regulated lncRNAs remained reduced until T16, while 40% were no longer regulated and 10% were even upregulated at T16 (**Figure 6C**). T8 showed the highest number of DEGs (2,531) with 1,766 exclusively regulated genes at this timepoint, while at T16, the number of DEGs was reduced again to 1,079, with 298 exclusively regulated genes (**Figure 6B**). This suggests that at longer SRSF5 depletion times, changes in gene expression are attenuated or overshoot due to compensatory mechanisms. In line with this, we identified very few DEGs upon siRNA-mediated SRSF5 KD (48 h) in P19 cells (Müller-McNicoll et al., 2016).

**Figure 6.**
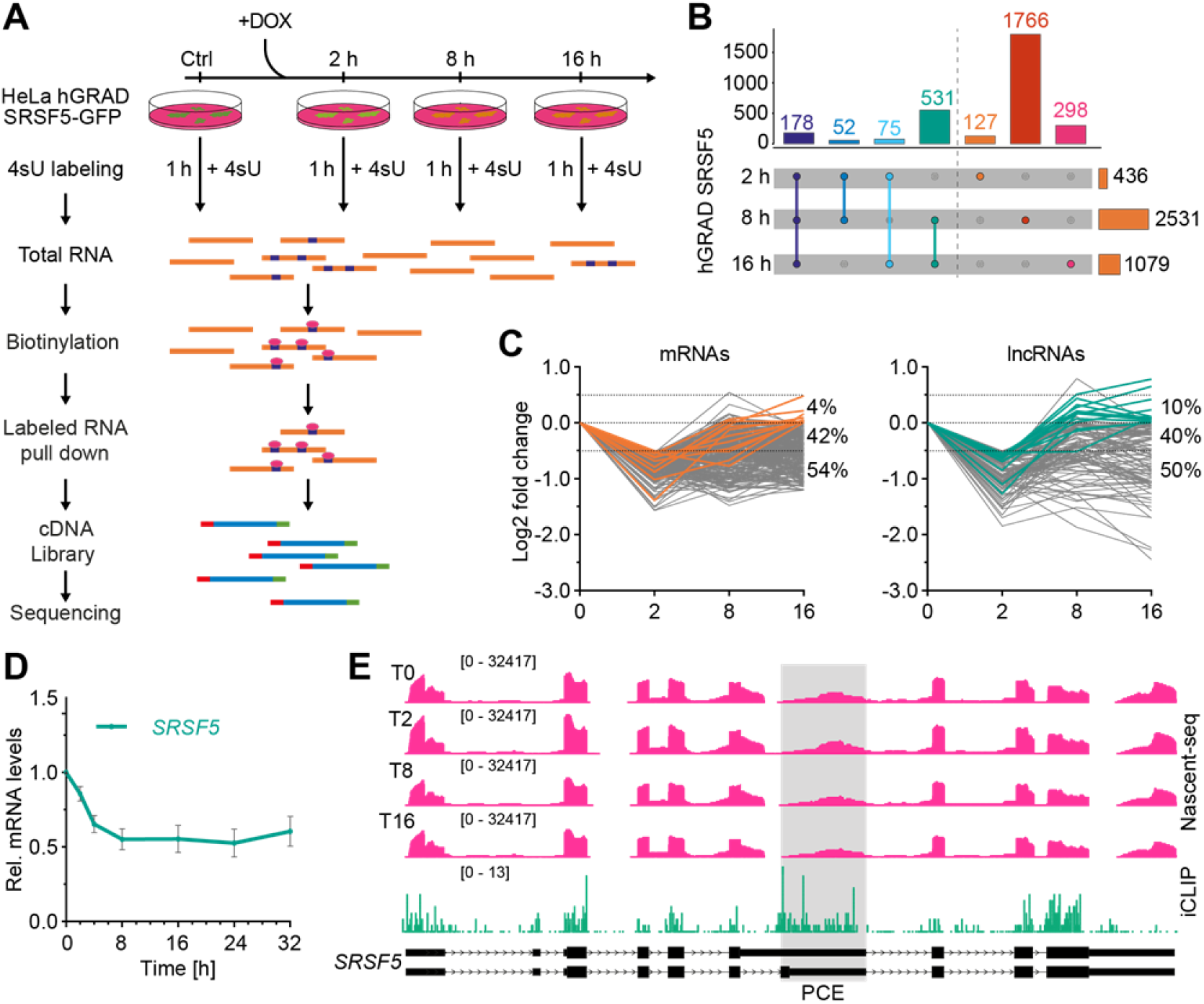
Combining hGRAD with Nascent-seq and iCLIP2 allows the identification of direct SRSF5 targets and reveals a function of SRSF5 in transcript stability. **A)** Scheme of the workflow combining a time-course of hGRAD degradation with Nascent-seq. Cells were first induced with DOX (1 μg/mL) for the indicated times. All new transcripts were labeled by addition of 4sU (400 μM) to the medium for one hour. Subsequently, total RNA was extracted, biotinylated, purified and converted into a cDNA library for deep sequencing. **B)** UpSet plot of differentially expressed transcripts. **C)** Rapid depletion of SRSF5 by hGRAD leads to dynamic changes in the abundance of mRNAs (left) and lncRNAs (right). All RNAs that are down- regulated at T2 are shown in grey. Transcripts that are compensated at T16 are highlighted in orange (mRNAs) or green (lncRNAs). **D)** RT-qPCR shows that *SRSF5* mRNA decreases until 8 h degradation and remains low until 32 h post-induction. Graph shows mean and SD of n=3 independent experiments. E) *SRSF5* transcripts show extensive SRSF5 binding but no processing alterations after SRSF5 depletion. Genome browser view showing Nascent-seq and SRSF5 iCLIP2 crosslinks on the *SRSF5* gene.

### SRSF5 binds preferentially to down-regulated transcripts

To test whether down-regulated transcripts are bound by SRSF5, we performed iCLIP2 (Buchbender et al., 2020) with HeLa hGRAD SRSF5-endo-GFP cells (5 replicates; **Figure S5E, S5F**). We identified 415,245 reproducible SRSF5 binding sites in 8,280 protein-coding genes, mainly in coding regions and 5ʹUTRs, and 708 lncRNAs (**Figure S5G, S5H**). Interestingly, 51% of the transcripts down-regulated at T2 contained SRSF5 binding sites compared to 15% in unregulated and 16.1% in up-regulated transcripts. A similar trend was observed for the other timepoints (**Figure S5J**), suggesting that those are direct targets of SRSF5. Among the bound transcripts, down-regulated transcripts tended to have more SRSF5 binding sites than up-regulated transcripts (0.77 vs 0.21; **Figure S5J**). For example, the lincRNAs *LINC01895*, *BASP1-AS, ENSG00000231412* and *ENSG00000259621* were extensively bound by SRSF5 and showed decreased levels at all three timepoints (**Figure S5I, S5K**). Similarly, *SRSF2* mRNA was decreased and highly bound by SRSF5 (**Figure S5K**).

Unexpectedly, another mRNA that was strongly bound by SRSF5 and downregulated at T8 and T16 was the *SRSF5* transcript itself. We expected its mRNA levels to rise upon depletion of the SRSF5 protein, since SRSF5 auto-regulates its own mRNA levels via splicing of a poison cassette exon (PCE), similar to other SR proteins (Müller-McNicoll et al., 2019). RT-qPCR confirmed that *SRSF5* transcript levels decreased to 50% and remained low even at 32 h of SRSF5 depletion (**Figure 6D**). SRSF5 bound most strongly in a region surrounding the *SRSF5* PCE, suggesting that SRSF5 binding normally regulates PCE inclusion (**Figure 6E**). In line with this, quantification of PCE junctions revealed a decrease in PCE inclusion after 8 and 16 h SRSF5 depletion from 10% to 4% (**Figure S6A**). This meant that the levels of *SRSF5-PCE* isoforms decreased, while those of translatable *SRSF5* isoforms increased at T16, which may counteract degradation of SRSF5. Apart from *SRSF2* and *SRSF5*, other SR protein encoding mRNAs did not change upon SRSF5 depletion. However, we observed an increase in PCE inclusion at T16 for SRSF3 (from 50.3 to 55.3%), SRSF6 (from 38.3 to 47.7%) and SRSF7 (from 63.5 to 69.7%; **Figures S6B-D**). Fewer translatable isoforms of these SR proteins may contribute to the observed compensatory effects at longer depletion times. Our data indicate that auto- and cross-regulation via PCE inclusion also operate when SR protein levels are too low.

But what causes the early decrease in the levels of *SRSF5* mRNA and other transcripts? SRSF5 could promote their transcription, splicing, 3ʹend processing and/or stability. Since the sudden decrease in *SRSF5* mRNA levels was also detectable with total RNA (**Figure 6D**), which is dominated by ʹoldʹ transcripts, reduced transcription could be ruled out. To test whether SRSF5 depletion caused intron retention (IR) or intronic polyadenylation, we analyzed the Nascent-seq data with IRFinder (Lorenzi et al., 2021). We identified 181 significant IR events at timepoint T2, 717 events at T8, and 142 events at T16 (**Table S1**). However, only 1.1% of the down-regulated transcripts showed IR at T2 (7.1% at T8, 1.4% at T16), and there was no evidence for premature polyadenylation. This suggests that IR is not the main cause of decreased transcript levels.

We speculated that *SRSF5-PCE* isoforms and other non-coding transcripts might decrease in abundance after acute SRSF5 depletion, because SRSF5 normally binds and targets them to nuclear speckles, which may protect them from nuclear decay machineries. To test this, we combined SRSF5 hGRAD depletion with RNA fluorescence *in situ* hybridization (FISH). We found that *SRSF5-PCE* transcripts accumulated in a bright spot that co-localized with SRSF5 and the nuclear speckle marker SRRM2 (**Figure S6E**). Interestingly, after 6 h of SRSF5 depletion *SRSF5-PCE* transcript levels were strongly decreased, the bright spots were completely dispersed and co-localization with nuclear speckles was no longer observed (**Figure S6E**). This suggest that acute SRSF5 degradation may release *SRSF5-PCE* transcripts from nuclear speckles and destabilize them, while normal SRSF5 isoforms are not affected and thus their relative levels increase. Based on the rapid decline of many other non-coding transcripts at 2 h after SRSF5 depletion, the preferential binding of SRSF5 to these transcripts and the absence of splicing alterations, we propose that SRSF5 binds and targets those transcripts to nuclear speckles to protect them from decay.

Our study demonstrates the usefulness of hGRAD to study the role of nuclear condensates in gene expression and discover novel functions of nuclear RBPs by distinguishing direct from indirect or compensatory effects.

## Discussion

Studying the functions of individual RBPs is a challenging task. This is due to similar domain structure, redundant functions, cooperation and competition for binding sites and large networks of auto- and cross-regulation (Leclair et al., 2020; Meinke et al., 2020; Sliskovic et al., 2022). Moreover, their high mRNA translation rates and protein stability require long depletion times when using RNAi, which causes indirect effects (Chen et al., 2018). To discriminate between the functions of individual SR proteins, we and others have used endogenous GFP-tagging by CRISPR/Cas9. Here, we introduce a simple system that allows the inducible, rapid and efficient depletion of such GFP-tagged RBPs through genomic integration of a single plasmid. We show that our system, termed hGRAD, can deplete various GFP-tagged proteins expressed in different cell types and localizing to different subcellular compartments, including nuclear condensates such as nuclear speckles and paraspeckles. hGRAD is more efficient, less toxic and requires less extensive genome editing than comparable approaches like the AID and Trim-Away systems.

hGRAD is derived from the deGradFP system, which was designed for degradation of GFP-tagged proteins in *Drosophila* (Caussinus et al., 2011). Several reports indicated that it is not suitable for the depletion of nuclear proteins in mammalian cells (Daniel et al., 2018; Li et al., 2019). To improve this, we exchanged the F-box domain from *Drosophila* Slmb for that of its human orthologue FBXW11. A similar approach named zGRAD was shown to strongly increase degradation efficiency in zebrafish (Yamaguchi et al., 2019). hGRAD achieved a similar degradation efficiency compared to deGradFP and a significantly faster depletion than zGRAD.

Compared to the auxin-inducible systems osTIR1 and aaAFB2 and to TRIM-away, hGRAD did not impair cell proliferation upon induction. This is likely because the other approaches require the constitutive expression of large (70-90 kDa) heterologous E3-Ub-ligases in order to be efficient (Clift et al., 2017; Daniel et al., 2018; Li et al., 2019), which contain additional enzymatic domains, making potential off-target effects more likely. The osTIR1 and aaAFB2 systems showed a faster onset of depletion with half-lives down to 25 min (Daniel et al., 2018), because all required protein components are already expressed prior to induction with auxin. Moreover, a fast recovery of the target protein can be achieved after auxin removal (Daniel et al., 2018; Li et al., 2019). hGRAD has a longer onset and does not allow target protein recovery similar to the deGradFP and zGRAD systems (Caussinus et al., 2011; Yamaguchi et al., 2019). This might be due to the high stability of the nanobody fusions and the single layer of induction via DOX addition. However, the osTIR1 system is intrinsically leaky and led to a significant degradation of SRSF3 prior to auxin addition, supporting other reports (Li et al., 2019). Moreover, the continuously expressed AID- nanobody possibly binds to GFP-tagged proteins and alters or inhibit their functions, which might contribute to the observed cytotoxic side effects.

Using hGRAD, we were able to almost completely deplete the nuclear speckle protein SRSF5 after 3 h, which otherwise requires long depletion times by RNAi. This allowed us to study its immediate effects with minimized compensation and cross-regulation. The combination of hGRAD with Nascent-seq uncovered highly dynamic transcript changes after SRSF5 depletion. Interestingly, these changes were transient in nature and already appeared much attenuated at 16 h, which illustrates the robustness of gene expression through compensatory mechanisms likely by other SR proteins. Indeed, we discovered that SRSF5 depletion alters the inclusion levels of its own PCEs as well as that of other SR proteins, indicating that auto- and cross-regulation operate when SRSF5 levels are too low. Moreover, our data suggest a novel role for SRSF5 in transcript stability, whereby SRSF5 binding targets and stores *SRSF5-PCE* transcripts and likely other non-coding transcripts in nuclear speckles. Acute depletion of SRSF5 releases bound transcripts from these condensates where they are now exposed to nuclear decay. At later time points other SR proteins might bind and compensate for nuclear speckle targeting so that transcript levels recover.

Rapid depletion is the tool of choice to study the functions of individual RBPs. hGRAD now allows the rapid degradation of notoriously difficult to target nuclear RBPs that localize to condensates. It allows to disentangle protein functions from those of transcripts that are produced from the same genomic locus, to study auto- and cross-regulation and it allows to study the role of condensates in the regulation of gene expression. Our ʹone-fits-allʹ system can be inserted by a one-step procedure in any cell expressing GFP-tagged proteins, often commercially available and functionally validated. Moreover, the hGRAD vector is flexible and the F-box domain could be replaced with other targeting moieties or enzymatic activities, e.g. APEX2 for proximity labelling or kinases to modify GFP-tagged proteins in a rapid and inducible manner, making it a very useful tool for future studies.

## Acknowledgements

We thank Maria Clara Hernandez Cañas for running IRFinder, Ina Poser (MPI-CBG) for the GFP-tagged BACs and Anke Busch and IMB Bioinformatics Core Facility for processing the iCLIP2 data. Support by the IMB Genomics Core Facility and the use of its NextSeq500 (funded by the Deutsche Forschungsgemeinschaft (DFG, German Research Foundation) – 329045328) is gratefully acknowledged. We are also grateful for funding from the Deutscher Akademischer Austauschdienst (DAAD) to IS and Deutsche Forschungsgemeinschaft (SFB902-B13 to BA, KZ and MMM).

## Author contributions

BA and MMM designed the experiments. BA, RJR, FM and EKO performed the experiments. BA, RJR, IS, MK and KZ performed the analyses. Figures were prepared by BA and RJR. The manuscript was written by BA, RR, KZ and MMM.

## Data availability

All sequencing data is available in the Gene Expression Omnibus (GEO) under the SuperSeries accession number GSE229326. The collection includes the Nascent-seq data of SRSF5-GFP-endo (GSE229324) as well as the iCLIP data for SRSF5-GFP-endo in HeLa cells (GSE229325). The token for anonymous reviewer access is ejsfsuaoxrkdfwh at: https://www.ncbi.nlm.nih.gov/geo/query/acc.cgi?acc=GSE229326

## Material and Methods

### Generation of plasmid constructs

The pTRE-BI-hGRAD, -TRIM21, -aaAFB2 and -osTIR1 plasmids were generated by Gibson assembly (GBA). For this a DOX-inducible bi-directional pTRE-BI vector was designed and synthesized by VectorBuilder (Vector ID: VB190904-1039fwc). For the pTRE-BI-hGRAD plasmid, a sequence encoding mCherry was added on one side of the tet-inducible bi-directional promoter by conventional restriction cloning (*Bam*HI & *Spe*I; New England Biolabs). The coding region of a human F-box domain (215 amino acids [aa], UniProt ID: Q9UKB1) fused to the sequence of an anti-GFP nanobody (VHH-GFP4) (Saerens et al., 2005) was inserted on the other side by GBA. For the pTRE-BI-TRIM21 plasmid, the pTRE-BI vector was linearized with *Bam*HI and *Nhe*I (New England Biolabs) and inserts were integrated by GBA. The sequences encoding TRIM21 (476 aa, UniProt ID: P19474) fused to mCherry and a weak NLS signal (AAAKRVKLD) (Li et al., 2019) was inserted at one side of the tet-inducible bi-directional promoter and the human IgG1 Fc2 domain was inserted at the other side. For pTRE-BI-osTIR1, the *osTIR1* gene (574 aa, UniProt ID: Q7XVM8) fused to mCherry and a weak NLS signal was inserted at one side of the bi-directional promoter and the mAID sequence (Daniel et al., 2018) and the GFP nanobody (VHH-GFP4) was added to the other side. For pTRE-BI- aaAFB2, the *aaAFB2* gene (575 aa, UniProt ID: Q9LW29) fused to mCherry and a weak NLS signal was inserted at one end of the bi-directional promoter and the miniIAA7 AID sequence (Li et al., 2019) and the GFP nanobody (VHH-GFP4) to the other side.

Plasmids used as circular homology directed repair (HDR) templates were generated by GBA. 1-kb homology arms flanking the single guide RNA (sgRNA) cut sites were amplified by PCR from host genomic DNA with primers containing overlapping sequences. The GFP-Neo^®^ resistance cassette containing a GFP-tag, the Neomycin (Neo^®^) resistance gene and an internal ribosome entry site (IRES) was amplified from the SRSF3 BAC DNA (Müller-McNicoll et al., 2016). PCRs were performed with either S7 Fusion Polymerase™ (Mobidiag), Q5^®^ High-Fidelity or Taq DNA Polymerases (both New England Biolabs). PCR inserts and linearized pGEM^®^-T Easy vector (Promega) were combined by GBA performed with Taq DNA Ligase and T5 Exonuclease (both New England Biolabs). One Shot™ TOP10 chemically competent *E. coli* cells (Invitrogen) were used for transformations. Plasmids were extracted from bacteria with the ZR Plasmid Miniprep Classic (Zymo Research). Successful cloning was confirmed by Sanger sequencing (ACGT). All plasmids are listed in **Table S4** and all primers in **Table S7**.

### Generation of stable cell lines and drug treatments

For integration of the pTRE-BI vectors, WT HeLa and P19 cells were transfected with 0.8 μg purified and linearized plasmid DNA per well in 6-well plates using the jetPRIME^®^ Transfection reagent (Polyplus-transfection). Cells with stably integrated plasmids were selected with 2 µg/mL puromycin (Gibco™, Thermo Fisher Scientific). Single cell clones were generated by limited dilution, grown from single cells in 96-well plates and screened for high inducibility by confocal microscopy upon DOX-induction (1 μg/mL).

Bacterial artificial chromosomes (BAC) harboring sequences encoding for C-terminally GFP-tagged *SRSF3, SRSF5, NONO, EIF4E* and *SRPK1* genes (**Table S5**) were isolated from *E. coli* DH10 cells using the NucleoBond™ Xtra Midi EF kit. For integration of the BACs WT HeLa and P19 cells were transfected with 1 μg purified BAC DNA per well in 6-well plates using jetPRIME®. Cells with stably integrated BACs were selected with 400 μg/mL Geneticin™ (G418, Gibco™, sorted for single cell clones with near-endogenous expression levels and expanded. All stable cell lines are listed in **Table S6**. HeLa and P19 cells were cultivated under humidified condition at 5% CO_2_ and 37 °C in DMEM GlutaMAX Medium, supplemented with 10% (v/v) heat inactivated fetal bovine serum and 100 μg/mL penicillin-streptomycin (all Gibco™, Thermo Fisher Scientific).

Live cells were counted with an EVE™ Automated Cell Counter device (NanoEnTek Inc). For this, equal volumes of cell suspension and 0.4% (v/v) trypan blue stain solution (Gibco™, Thermo Fisher Scientific) were mixed and applied to EVE™ Cell counting slides. To inhibit the proteasome, HeLa cells were grown on coverslips placed in 24-well plates and treated with 10 μM of MG132 (M7449-200UL, Sigma Aldrich) diluted in fresh DMEM for 4 h followed by fixation with 4% PFA (Thermo Fisher Scientific). To induce expression of the hGRAD and TRIM21 systems, cells were treated with 1 μg/mL doxycyclin (DOX; Sigma-Aldrich, D9891). osTIR1 and aaAFB2-expressing cells were additionally treated with 0.5 mM auxin (AUX; Sigma Aldrich, I2886-25G).

### Genome editing with CRISPR/Cas9

For endogenous C-terminal tagging of *SRSF3*, *SRSF5* and *SRRM2*, single guide RNAs (sgRNAs) were designed using CRISPOR (http://crispor.tefor.net) and purchased from IDT (Integrated DNA Technologies). HeLa and P19 cells were first transfected with circular HDR donor plasmids comprising the GFP-Neo^®^ resistance cassette flanked by gene-specific 1-kb homology arms (HA) using jetPRIME^®^ (Polyplus). After 6-12 h, WT cells were transfected with pre-assembled sgRNAs (Alt-R^®^ CRISPR-Cas9 crRNA (target-specific, see **Table S2**) and Alt-R^®^ CRISPR-Cas9 tracrRNA Atto550 labeled, IDT) and recombinant Cas9 protein (Alt-R^®^ *S. p.* HiFi Cas9 Nuclease V3, IDT) using Lipofectamine™ CRISPRMAX (Invitrogen) and cultured for 48 h in the presence of an HDR enhancer (final concentration 20 μM, IDT). Cells were then selected with Geneticin^®^ (Gibco, 400 μg/mL). CRISPR clones were generated by limited dilution and grown from single cells in 96-well plates. For genomic screening, cells were washed twice with PBS in the 96-well plates and lysed in directPCR buffer (20 mM Tris-HCl, pH 8, 200 mM NaCl, 1 mM EDTA, 0.5% Tween-20, 0.5% NP-40) with freshly added 200 μg/mL Proteinase K (Sigma Aldrich), and incubated for 1 h at 55°C followed by Proteinase K inactivation at 95 °C for 15 min. Screening PCRs were performed from crude lysates using primers flanking the edited region. Sequences of the gRNAs are shown in **Table S2** and primers in **Table S7.**

### RNA isolation, reverse transcription and qPCR

Cells were harvested and resuspended in TRIzol™ (Thermo Fisher Scientific). RNA was extracted according to the manufacturer’s instructions, treated with TURBO™ DNase (Thermo Fisher Scientific) for 30 min at 37 °C to remove genomic DNA and subsequently purified. Two μg of RNA were reverse transcribed into cDNA using SuperScript™ and 10 mM dNTP Mix (both Thermo Fisher Scientific) and oligodT (Sigma Aldrich). qPCR primers were selected using Primer-BLAST (https://www.ncbi.nlm.nih.gov/tools/primer-blast/). qPCRs were performed using cDNA (1:8 dilution) and the ORA™ SEE qPCR Green ROX L kit (highQu) on a PikoReal 96 machine (Thermo Fisher Scientific). GraphPad Prism was used for graphics/statistics. Primers used are listed in **Table S7**.

### Western blot and antibodies

Cells were lysed in 300 μl NET-2 buffer (150 mM NaCl, 0.05% (v/v) NP-40, 50 mM Tris-HCl pH 7.5), supplemented with 1× cOmplete Protease Inhibitor Cocktail (Sigma Aldrich) and 10 mM β-phosphoglycerate (Fluka BioChemica) or with RIPA buffer (150 mM NaCl, 0.05% (v/v) NP-40, 50 mM Tris-HCl pH 7.5, 0.1% (w/v) SDS, 0.5% (w/v) sodium deoxycholate, freshly added 1× cOmplete Protease Inhibitor Cocktail and 10 mM β-phosphoglycerate). NET-2 lysates were sonicated on ice for 30 sec (3 pulses of 10 sec; 20 sec intervals) at 20% amplitude (Branson W-450 D) and cleared by centrifugation. Protein concentrations were measured using Quick Start Bradford 1× Dye Reagent (Bio- Rad^®^) on a NanoDrop2000 (Thermo Fisher Scientific) or DC Protein-Assay (Bio-Rad^®^) for RIPA samples. 20-40 μg protein were separated by SDS-PAGE on 4-15% Mini- PROTEAN^®^ TGX Stain-Free™ Gels (Bio-Rad^®^). and transferred onto PVDF membranes using Trans-Blot Turbo RTA Mini LF PVDF Transfer Kit (Bio-Rad^®^). Transfer and equal loading were evaluated by activation of stain-free gels by UV light. Membranes were probed with the antibodies listed in **Table S3**. Proteins were imaged using secondary antibodies coupled to a horseradish peroxidase and Amersham™ ECL Prime Western Blotting Detection Reagent (Cytiva) with the ChemiDoc™ MP Imaging System. Image quantification was performed using the ImageLab software (Bio-Rad).

### GFP fluorescence microscopy and live-cell imaging

For GFP fluorescence microscopy, cells were grown on 10 mm glass coverslips. After removing the medium cells were gently washed twice with DPBS and fixed with 4% PFA (Thermo Fisher Scientific) in PBS for 10 min at RT. DNA was stained with Hoechst 34580 (1:4000, in TBST; Thermo Fisher Scientific). After a final wash, the coverslips were dried and mounted on ProLong™ Diamond Antifade Mountant (Invitrogen). All microscopy experiments were visualized using a Zeiss LSM780 confocal microscope system and Zen Black imaging software.

For live-cell imaging, cells were incubated in a microscopy grade 24-well glass-bottom culture plate (Sensoplate, Greiner Bio-ONE). Prior to imaging, the medium was exchanged with low autofluorescent FluoroBrite™ DMEM Media (Gibco) supplemented with 10% (v/v) FBS, 2 mM GlutaMAX ™ I Supplement and 1 mM sodium pyruvate (all Gibco™, Thermo Fisher Scientific). 100 nM SiR-DNA stain (Spirochrome) was added for the visualization of nuclei. Cell culture plates were mounted on Zeiss LSM 780 microscope, equipped with a humidified incubation chamber (pre-heated for 2 hours at 37 °C, 5% CO_2_). Selected regions were automatically imaged every 15-20 minutes with pre-set settings and software autofocus using the ZEN Black software. GFP intensity and SiR-DNA were visualized in parallel. Eight Z-stacks per region with a slice thickness of 2 μm were imaged. To minimize phototoxicity, minimal possible laser intensity was chosen (488 nm <3%, 670; 633 nm <5%). GFP intensity changes in individual cells were quantified over time using Fiji software (Schindelin et al., 2012) and the TrackMate plugin (Tinevez et al., 2017). First, a Z-maximum intensity projection was made to generate a single plane image. The background was subtracted using a rolling ball background subtraction algorithm with a disk size of 75 μm. TrackMate was first applied to the SiR-DNA channel to identify and connect cell tracks over time. The mean GFP intensity was evaluated in corresponding cells and normalized to the first time point.

### 4sU treatment and nascent RNA sequencing

HeLa cells were treated with 400 μM 4-thio-uridine (4sU, Sigma Aldrich) for 1 hour to label freshly transcribed RNAs. The reaction was stopped by applying TRIzol™RI reagent to the culture dishes. RNA extraction and pulldown of newly transcribed RNA was performed according to (Gressel et al., 2019) without fractionation of the RNA by sonification. Nascent RNA was purified using RNA Clean and Concentrate Kit (ZymoResearch). cDNA libraries for RNA-seq were prepared with universal Plus™ Total RNA-Seq Library Preparation Kit (Tecan) according to manufacturerʹs instructions. Ribosomal RNA fragments were removed and the library was sequenced on an Illumina NovaSeq6000 instrument (2 replicates per timepoint, 100 Mio reads, 150 bp, paired-end). Reads were mapped to the human genome (hg38) using the STAR mapper (version 2.7.10) (Dobin et al., 2013) with the following parameters: --outFilterMultimapNmax 1 -- outFilterMismatchNmax 999 --outFilterMismatchNoverReadLmax 0.04 --outSAMtype BAM SortedByCoordinate.

To quantify IR events, IRFinder (version 1.3.1) (Middleton et al., 2017) was used with default parameters. Differential gene expression was quantified using DESeq2 with default parameters (Love et al., 2014). Sashimi plots for poison cassette exons of selected SR proteins were prepared using the R/Bioconductor package Gviz (version 1.37.2) by taking the merged BAM files of each timepoint as input. Junctions were filtered for the junctions describing the inclusion levels of the poison cassette exons.

### iCLIP2 libraries

iCLIP experiments were performed using the iCLIP2 protocol (Buchbender et al., 2020) with minor modifications. For each replicate, cells were grown near confluence on two 150-mm culture dishes, washed with ice-cold PBS, irradiated with 150 mJ/cm^2^ UV light at 254 nm (CL-1000, UVP), harvested by scraping and centrifugation and stored at -80°C until lysis. Following lysis and partial digestion with RNase I (Thermo Fisher Scientific, AM2294), immunoprecipitation of SRSF5-GFP was performed using a goat anti-GFP antibody (MPI-CBG, Dresden, Germany) coupled to Dynabeads™ Protein G (Thermo Fisher Scientific, 10002D). Co-purified, crosslinked RNA fragments were dephosphorylated at their 3’ ends using T4 Polynucleotide Kinase (New England Biolabs, M0201S) and ligated to a pre-adenylated 3’ adapter (L3-App). To visualize protein-RNA complexes, RNA fragments crosslinked to SRSF5-GFP were labeled at their 5’ ends using T4 Polynucleotide Kinase and γ-^32^P ATP (Hartmann Analytic). Samples were run on a Nu-PAGE 4-12% Bis-Tris Protein Gel (Thermo Fisher Scientific, NP0335BOX), transferred to a 0.45 μm nitrocellulose membrane (GE Healthcare Life Science, 10600002) and visualized using a Phosphorimager. Regions of interest were cut from the nitrocellulose membrane (70 kDa to 130 kDa) and RNA was released from the membrane using Proteinase K (Roche, 03115828001). RNA was purified using neutral phenol/chloroform/isoamylalcohol (Ambion, AM9722) followed by chloroform (Serva, 39554.02) extraction, and reverse transcribed using SuperScript III™ (Life Technologies, 18080-044). cDNA was cleaned up using MyONE Silane beads (Life Technologies, 37002D) followed by ligation of a second adapter containing a bipartite (5-nt + 4-nt) unique molecular identifier (UMI) as well as a 6-nt experimental barcode (Buchbender et al., 2020). iCLIP2 libraries were pre-amplified with 6 PCR cycles using short primers (P5Solexa_short and P3Solexa_short) and then size-selected using the ProNex Size-Selective Purification System (Promega, NG2001) in a 1:2.95 (v/v) sample:bead ratio to eliminate products originating from short cDNAs or primer dimers. The size-selected library was amplified for 6 cycles using P5Solexa and P3Solexa primers, and primers were removed using the ProNex Size-Selective Purification System (Promega, NG2001) in a 1:2.4 (v/v) sample:bead ratio. Purified iCLIP2 libraries were sequenced on a NextSeq500 System (Illumina) using a NextSeq^®^ 500/550 High Output Kit v2 as 92-nt single-end reads including a 6-nt sample barcode as well as 5+4-nt UMIs yielding between 5.4 and 74.1 million reads.

### iCLIP2 analysis

Basic quality controls were done using FastQC (version 0.11.8) (https://www.bioinformatics.babraham.ac.uk/projects/fastqc/). The FASTX-Toolkit (version 0.0.14) (http://hannonlab.cshl.edu/fastx_toolkit/) and seqtk (version 1.3) (https://github.com/lh3/seqtk/) were used to filter reads based on sequencing qualities (Phred score) in the barcode and UMI regions. Read were de-multiplexed according to the sample barcode on positions 6 to 11 of the reads using Flexbar (version 3.4.0, using non-default parameter --barcode-keep) (Roehr et al., 2017). Flexbar was also used to trim UMI and barcode regions as well as adapter sequences from read ends requiring a minimal overlap of 1 nt of read and adapter. UMIs were added to the read names and reads shorter than 15 nt were removed from further analysis. The downstream analysis was done as described in Chapters 3.4 and 4 of (Busch et al., 2020) with an additional step to remove reads directly mapped to the chromosome ends using Samtools (version 1.9) (Danecek et al., 2021) and bedtools (version 2.27.1) (Quinlan and Hall, 2010). Those reads do not have an upstream position and, thus, no crosslink position can be extracted. Genome assembly and annotation of GENCODE (release 31) (Frankish et al., 2019) were used during mapping with STAR (version 2.7.3a) (Dobin et al., 2013).

Processed reads from five replicates were merged prior to peak calling with PureCLIP (version 1.3.1) (Krakau et al., 2017) using a minimum transition probability of 1%. Significant crosslink sites (1 nt) were filtered by their PureCLIP score, removing the lowest 2% of crosslink sites. The remaining sites were merged into 7-nt wide binding sites using the R/Bioconductor package BindingSiteFinder (version 1.0.0), filtering for sites with at least 3 positions covered by crosslink events. Binding sites were overlapped with gene and transcript annotations obtained from GENCODE (release 29). Binding sites within protein-coding genes were assigned to the transcript regions, i.e., intron, CDS, 3ʹUTR or 5ʹUTR.

