## Supplement for "hGRAD – a versatile ʹone-fits-allʹ system for the acute depletion of RNA binding proteins in nuclear condensates"

### SUPPLEMENTARY MATERIAL

Content:

Supplementary Figures .......... 2
Supplementary Tables .......... 13

#### Supplemental Figures

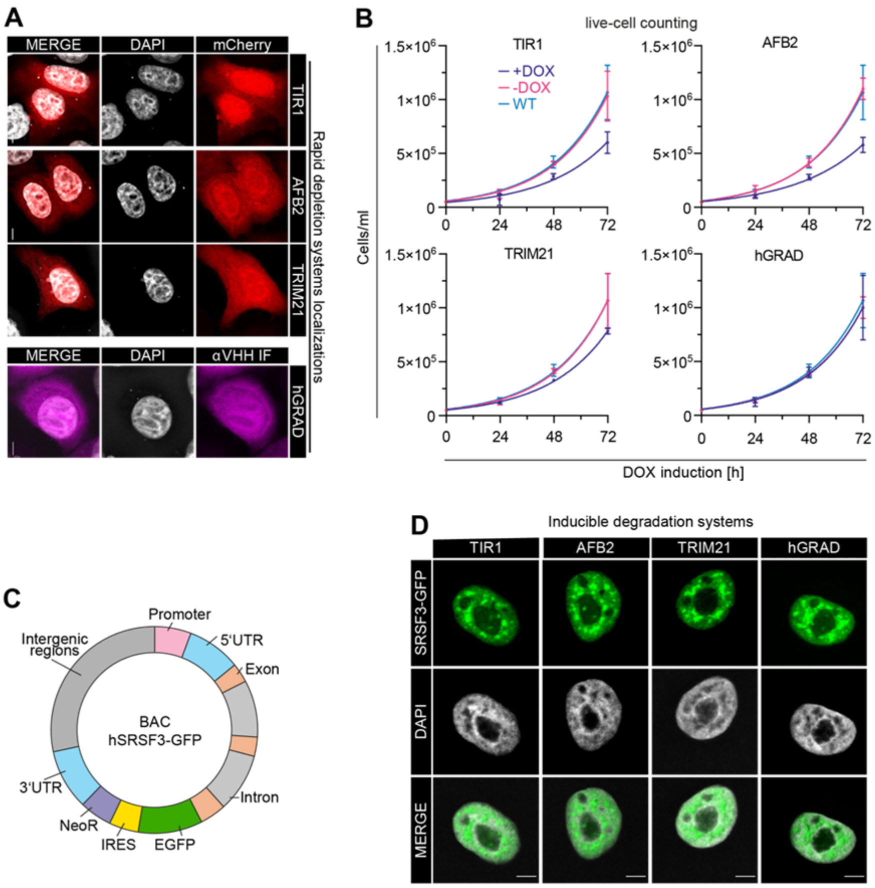


**Figure S1. hGRAD performs superior to other inducible degradation systems. A)** Subcellular localization of the induced proteins. Microscopy images of different HeLa master cell lines induced with DOX (1 µg/mL) for 16 h. **Upper panel:** TIR1-, AFB2-, and TRIM21-mCherry (red), DNA (grey). **Lower panel:** hGRAD F-box-nanobody fusion was detected with an anti-VHH antibody and an Alexa Fluor 647 dye coupled secondary antibody (magenta), DNA (grey). Scale bars = 5 µm. **B)** Experimental scheme and growth curves comparison of HeLa live cells expressing the hGRAD, TRIM21, osTIR1 or aaAFB2 systems until 72 h after DOX induction. The doubling time ($t_d$) was evaluated by exponential growth equation fit ($Y=Y_0*exp(k*X)$). WT control (blue), uninduced control (pink), induced (purple). Mean and standard deviation of the mean (SD) are shown from n=3 independent experiments. **C)** Scheme of a bacterial artificial chromosome (BAC) that was randomly integrated into the genome of HeLa cells. The BAC carries the complete human *SRSF3* gene fused to *EGFP* including the *SRSF3* promoter, 5'UTR, exons, introns and 3'UTR to ensure near endogenous expression and isoform levels. IRES - internal ribosome entry site, NeoR - Neomycin resistance cassette. **D)** SRSF3-GFP is expressed at similar levels in all degradation systems and localizes to nuclear speckles and the nucleoplasm in HeLa cells. Scale bars = 5 µm.

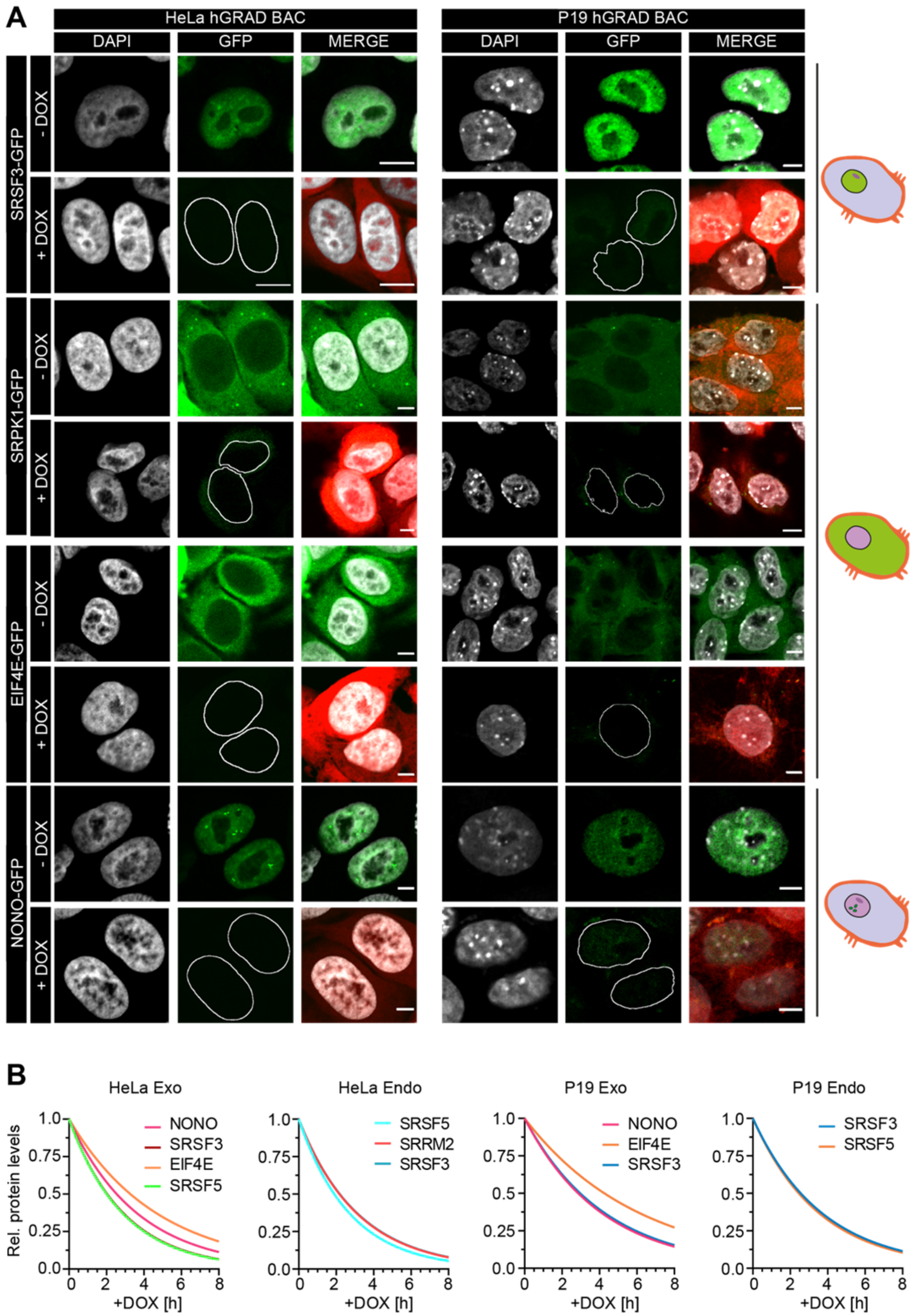


**Figure S2. hGRAD works in different cell types and species. A)** Representative micrographs showing the subcellular localization and degradation of GFP-tagged SRSF3, SRPK1, EIF4E and NONO in human HeLa and mouse P19 cells 16 h after induction by DOX (1 μg/mL). GFP-tagged proteins were expressed from integrated BACs. Red channel shows mCherry induction control. Scale bars = 5 μm. **B)** Non-linear regression fit (Exponential One Phase Decay Model) to determine the protein half-lives of GFP-tagged proteins in HeLa and P19 cells based on Western blots.

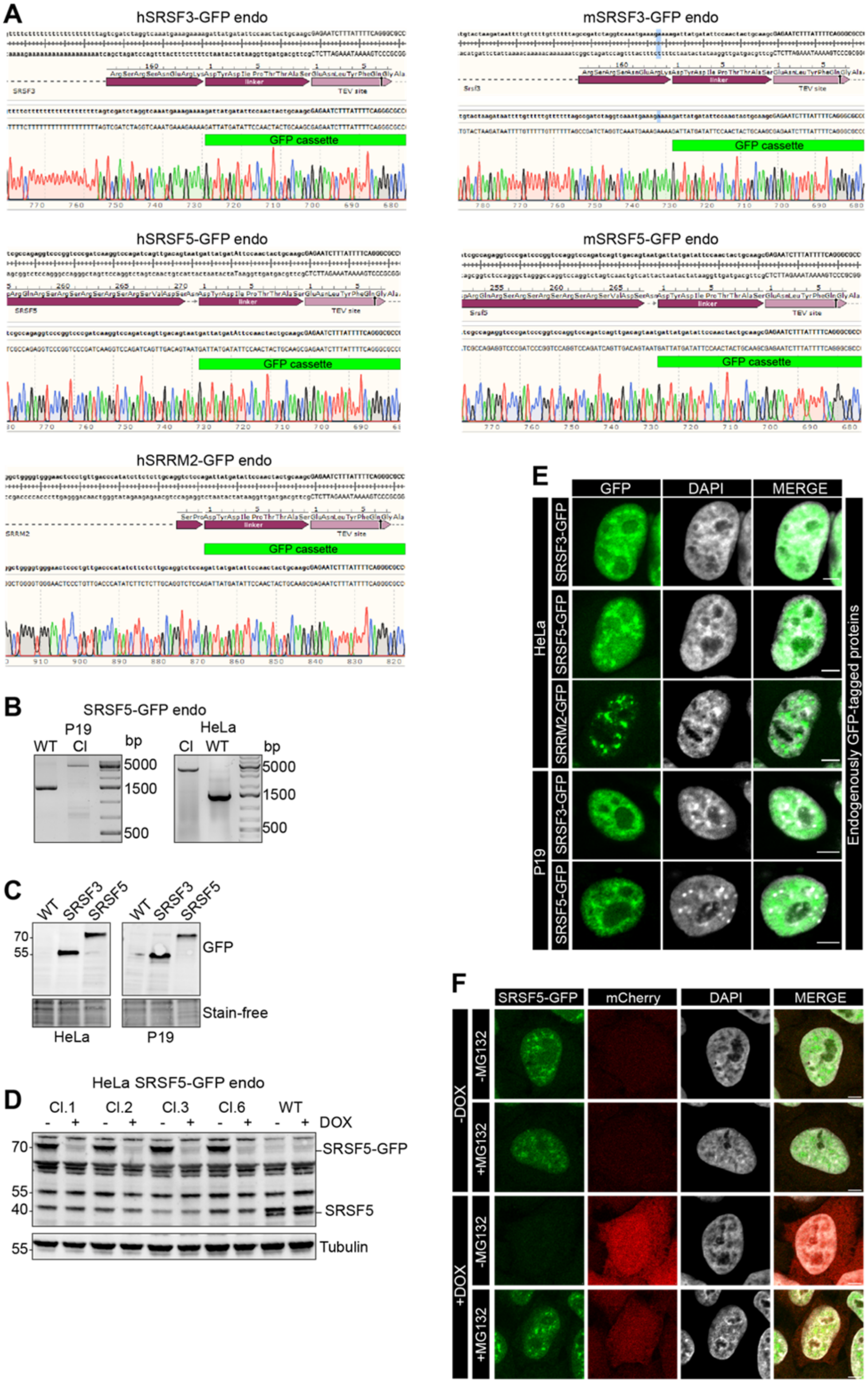


**Figure S3. hGRAD efficiently degrades endogenously GFP-tagged nuclear RBPs. A-B)** Correct insertion of the GFP cassette in the genomic locus of human (hSRSF3, hSRSF5 and hSRRM2) and mouse (mSRSF3 and mSRSF5) RBP genes was validated **A)** by Sanger sequencing, **B)** by PCR amplification of the genomic locus, **C)** by Western blot using an anti-GFP antibody and **D)** by Western blot using the anti-SRp40 (Sigma) antibody. Tubulin served as loading control. **E)** Subcellular localization of GFP-tagged proteins was validated by confocal microscopy. **F)** SRSF5 degradation is blocked when the proteasome is inhibited by MG132 for 4 h. All: Scale bars = 5μm.

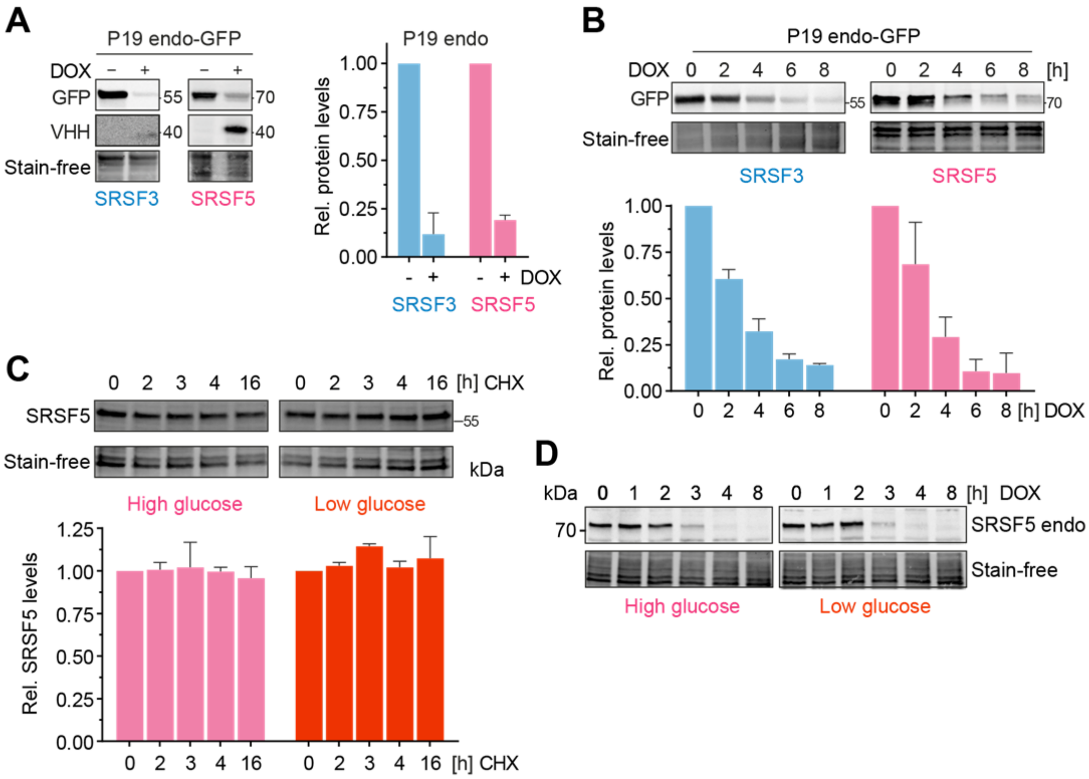


**Figure S4. hGRAD efficiently degrades endogenously GFP-tagged nuclear RBPs in P19 cells allowing their rapid knockdown. A)** Degradation efficiency of SRSF3- and SRSF5-endo-GFP in P19 cells 16 h after DOX induction (1 µg/mL). **B)** Degradation time-courses show that hGRAD works efficiently for SRSF3- and SRSF5-endo-GFP in P19 cells. **C)** Representative Western blot for a time course under high (4.5 g/L) and low (1.0 g/L) glucose conditions over 16 h upon treatment with the eukaryotic translation inhibitor cycloheximide (CHX; 10 µg/mL) to determine the stability of SRSF5-endo-GFP in HeLa cells. **A-C)** Shown are one representative Western blot and the quantification (mean and SD) of n=3 independent experiments. Quantifications relative to the -DOX or 0 h time point. Stain-free membranes were used to control for equal loading. **D)** Comparison of SRSF5-endo-GFP degradation by hGRAD in HeLa cells in high and low glucose conditions.

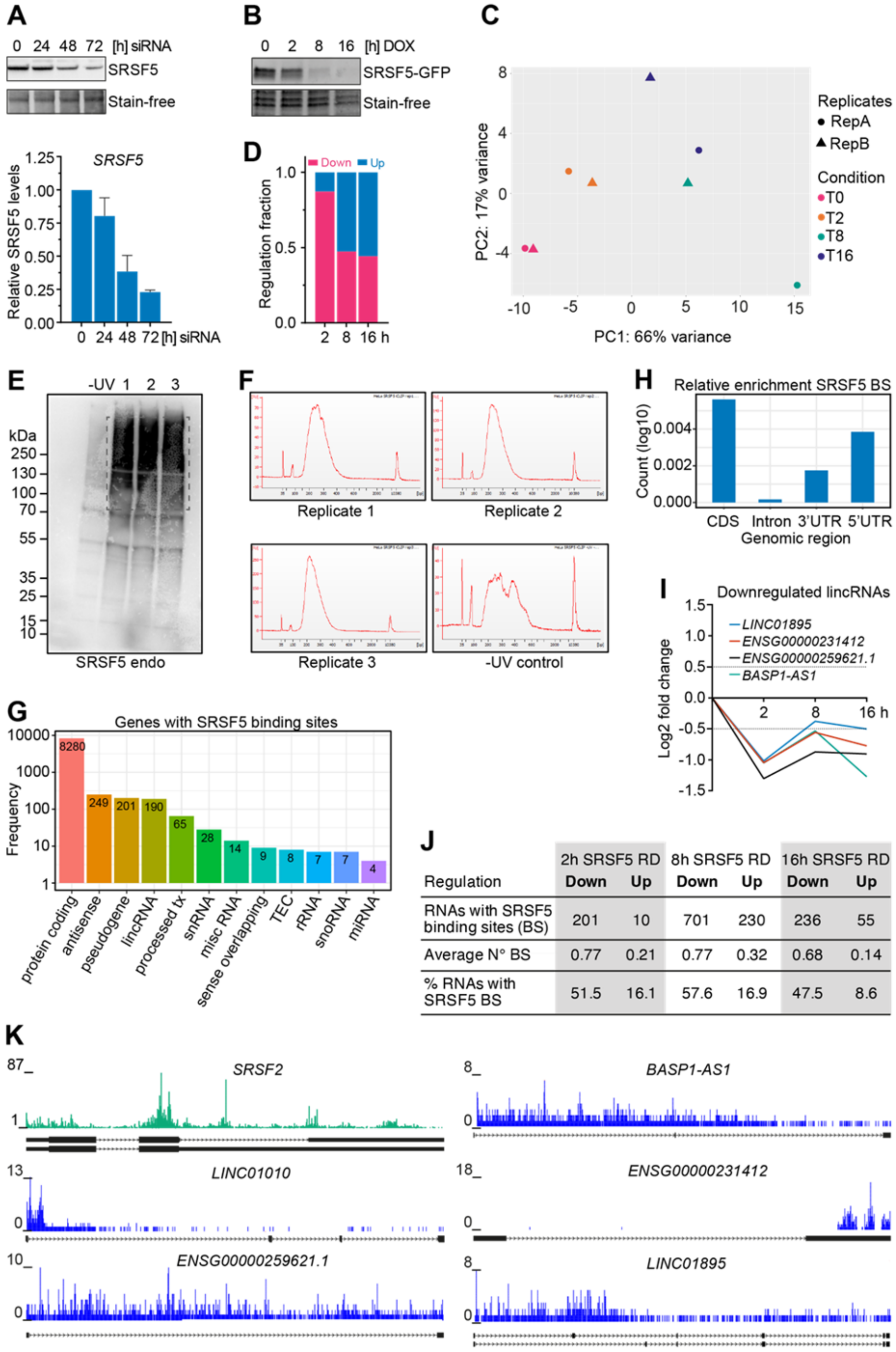


**Figure S5. Combining hGRAD with Nascent-seq allows the identification of direct SRSF5 targets. A) Top:** Representative Western blot for SRSF5 KD using siRNAs over a 72 h time-course experiment. **Bottom:** Quantification (mean and SD) relative to the 0 h sample of n=2 independent experiments. **B)** Representative degradation time-course of SRSF5-endo-GFP used for Nascent-seq. **C)** Principal component analysis showing the clustering of replicates in principal components (PC) 1 and 2. **D)** Fraction of down- and up-regulated transcripts at the T2, T8 and T16 time points after DOX induction. **E)** Autoradiograph of SRSF5-endo-GFP cells used for iCLIP2 (n=3 replicates). Non-crosslinked cells (-UV) were used as control. **F)** iCLIP2 libraries quantified using a Bioanalyzer instrument. **G)** Genes with SRSF5 binding sites sorted by RNA biotype. **H)** Enrichment of SRSF5 binding sites in specific transcript regions, i.e., coding region (CDS), introns, 3'UTR and 5'UTR relative to feature length. **I)** Examples for the dynamics of down-regulated lincRNAs. **J)** SRSF5 binding sites in regulated transcripts. **K)** Genome browser views showing SRSF5 crosslink events on *SRSF2* mRNA and selected lincRNAs that are down-regulated after rapid SRSF5 depletion.

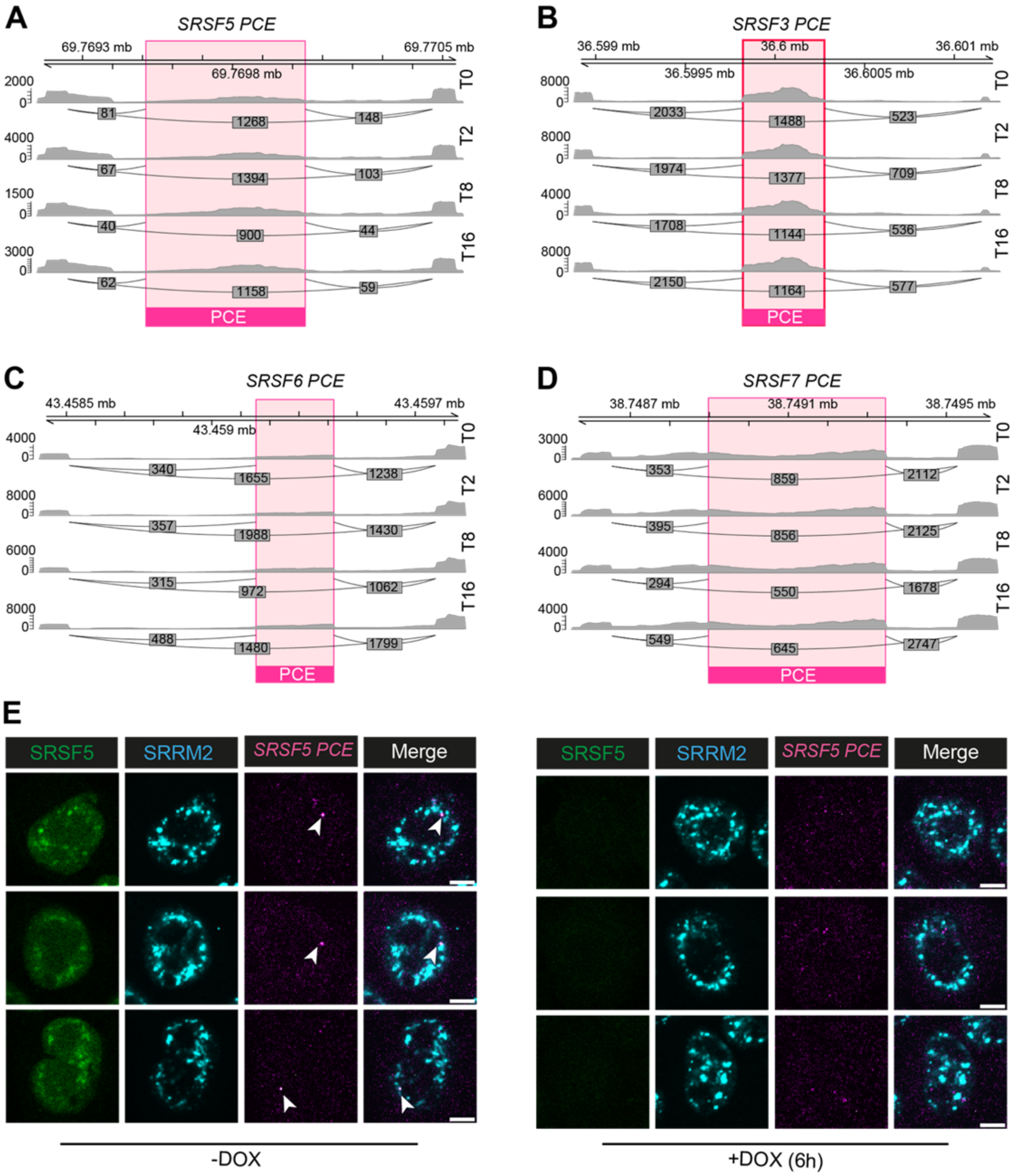


**Figure S6. *SRSF5-PCE* isoforms are reduced and dispersed after rapid depletion of SRSF5. A-D)** Sashimi plots with junction counts for skipped or included poison cassette exons (PCEs) of **A)** *SRSF5*, **B)** *SRSF3*, **C)** *SRSF6* and **D)** *SRSF7*. Only selected junctions are shown. **E) Left:** Representative micrographs showing the subcellular localization of SRSF5-endo-GFP and *SRSF5-PCE* isoforms in HeLa cells before DOX induction. **Right:** Representative micrographs showing the degradation of SRSF5-endo-GFP and the dispersal of *SRSF5-PCE* isoforms in HeLa cells after 6 h induction by DOX (1 μg/mL). Nuclear speckles were labelled with α-SRRM2 antibodies. *SRSF5-PCE* isoforms were labeled by RNA FISH. Scale bars = 5 μm.

#### Supplementary Tables

**Table S1: Integration of DESeq2, IRFinder and iCLIP2.**

**Table S2: List of sgRNAs.**

| Species | Name/Target | Protospacer sequence | PAM | Purpose | Supplier |
|---|---|---|---|---|---|
| Human | SRSF3 crRNA | gtcgatctaggtcaaatgaa | AGG | HDR, C-term. GFP-tag | IDT |
| Human | SRSF5 crRNA | aggtccagatcagttgacag | TGG | HDR, C-term. GFP-tag | IDT |
| Human | SRRM2 crRNA | ttccaccacacccaatgctc | TGG | HDR, C-term. GFP-tag | IDT |
| Mouse | SRSF3 crRNA | gccgatctaggtcaaatgaa | AGG | HDR, C-term. GFP-tag | IDT |
| Mouse | SRSF5 crRNA | aggtccagatcagttgacag | TGG | HDR, C-term. GFP-tag | IDT |
| - | universal tracrRNA | - | - | - | IDT |

**Table S3: List of antibodies used in this study.**

| Name | Species | Supplier | Catalog-Nr. |
|---|---|---|---|
| **α-Camelid-VHH** | rabbit | Genscript | A01860 |
| **α-GFP** | goat | Eric Geertsma, MPI-CBG | - |
| **α-mCherry** | rabbit | Abcam | ab167453 |
| **α-SRp40 (SRSF5)** | rabbit | Merck Millipore | 06-1365 |
| **α-goat-HRP** | donkey | Sigma Aldrich | AB324P |
| **α-rabbit-HRP** | donkey | Merck Millipore | AP182P |
| **α-rabbit-Alexa Fluor 680** | donkey | Thermo Fisher Scientific | A10043 |
| **α-rabbit-Alexa Fluor Plus 405** | donkey | Thermo Fisher Scientific | A48258 |
| **α-SRRM2** | rabbit | Thermo Fisher Scientific | PA5-59559 |

**Table S4: List of plasmids used or generated in this study.**

| Name | Vendor | Catalog Nr. | Reference |
|---|---|---|---|
| **pmCherry-N1** | Clontech | 632523 | - |
| **pGEM®-T Easy** | Promega | A1360 | - |
| **pTRE-BI** | VectorBuilder | VB190904-1039fwc | This work |
| **pSH-EFIRES-P-AtAFB2-mCherry-weak NLS** | Addgene | 129717 | (Li et al., 2019) |
| **pUAST_NSlmb-vhhGFP4** | Addgene | 35575 | (Caussinus et al., 2011) |
| **pCAGGs-NLS-TIR1_P2A_NES-TIR1** | Addgene | 117699 | (Daniel et al., 2018) |
| **pIRES_IBB-mCherry-IRES-HA-mAID-nanobody** | Addgene | 117720 | (Li et al., 2019) |
| **pSH-EFIRES-P-Seipin-miniIAA7-3XFlag** | Addgene | 129722 | (Li et al., 2019) |
| **pFUSE-hIgG1-Fc2** | Invivogen | #pfuse-hg1fc2 | - |
| **pTRE-BI-hGRAD-mCherry** | - | - | This work |
| **pTRE-BI-TRIM21-mCherry-hIgG1FC2** | - | - | This work |
| **pTRE-BI-TIR1-mCherry-mAID** | - | - | This work |

| Name | Vendor | Catalog Nr. | Reference |
|---|---|---|---|
| **pTRE-BI-AFB2-mCherry-miniIAA7** | - | - | This work |
| **pGEMT-HDR-hSRSF3-GFP** | - | - | This work |
| **pGEMT-HDR-mSRSF3-GFP** | - | - | This work |
| **pGEMT-HDR-hSRSF5-GFP** | - | - | This work |
| **pGEMT-HDR-mSRSF5-GFP** | - | - | This work |
| **pGEMT-HDR-hSRRM2-GFP** | - | - | This work |

**Table S5: List of BACs used in this study.**

| Internal-Nr. | Name | Selection marker |
|---|---|---|
| **MCB#2951** | mSRSF5-GFP | Geneticin / Chloramphenicol |
| **MCB#6553** | hSRSF5-GFP | Geneticin / Chloramphenicol |
| **MCB#5415** | mSRSF3-GFP | Geneticin / Chloramphenicol |
| **MCB#3204** | mSRPK1-GFP | Geneticin / Chloramphenicol |
| **MCB#5404** | mNONO-GFP | Geneticin / Chloramphenicol |
| **MCB#8367** | mEIF4E-GFP | Geneticin / Chloramphenicol |

**Table S6: List of cell lines used or generated in this study (selection marker: puromycin (Puro) or Geneticin (Gen) as indicated)**

| Name | Selection marker | Source/Reference | Species |
|---|---|---|---|
| **HeLa wild type** | - | ATTC® | Human |
| **HeLa hGRAD** | Puro | This work | Human |
| **HeLa TRIM21** | Puro | This work | Human |
| **HeLa TIR1** | Puro | This work | Human |
| **HeLa AFB2** | Puro | This work | Human |
| **HeLa hGRAD SRSF3-GFP (BAC)** | Puro / Gen | This work | Human |
| **HeLa TRIM21 SRSF3-GFP (BAC)** | Puro / Gen | This work | Human |
| **HeLa TIR1 SRSF3-GFP (BAC)** | Puro / Gen | This work | Human |
| **HeLa AFB2 SRSF3-GFP (BAC)** | Puro / Gen | This work | Human |
| **HeLa hGRAD SRSF5-GFP (BAC)** | Puro / Gen | This work | Human |
| **HeLa hGRAD SRPK1-GFP (BAC)** | Puro / Gen | This work | Human |
| **HeLa hGRAD EIF4E-GFP (BAC)** | Puro / Gen | This work | Human |
| **HeLa hGRAD NONO-GFP (BAC)** | Puro / Gen | This work | Human |
| **HeLa hGRAD SRSF3-GFP (HDR)** | Puro / Gen | This work | Human |
| **HeLa hGRAD SRSF5-GFP (HDR)** | Puro / Gen | This work | Human |
| **HeLa hGRAD SRRM2-GFP (HDR)** | Puro / Gen | This work | Human |
| **P19 wild type** | - | Sigma Aldrich / ECACC | Mouse |
| **P19 hGRAD** | Puro | This work | Mouse |
| **P19 hGRAD SRSF3-GFP (BAC)** | Puro / Gen | This work | Mouse |
| **P19 hGRAD SRSF5-GFP (BAC)** | Puro / Gen | This work | Mouse |
| **P19 hGRAD SRPK1-GFP (BAC)** | Puro / Gen | This work | Mouse |
| **P19 hGRAD EIF4E-GFP (BAC)** | Puro / Gen | This work | Mouse |
| **P19 hGRAD SRSF3-GFP (HDR)** | Puro / Gen | This work | Mouse |
| **P19 hGRAD SRSF5-GFP (HDR)** | Puro / Gen | This work | Mouse |

**Table S7: List of primers used in this study.**

Primers for cloning of rapid depletion vectors

| Primer | Primer Sequence (5'-3') | Insert / Amplicon |
|---|---|---|
| GBA Bi hGRADE ins1 f | ctttcttgtacaaagtgggctagcatggagcccgactcg | Insert 1 hFBXW11 |
| GBA Bi hGRADE ins1 r | agttggacgtgtcgtccaca | Insert 1 hFBXW11 |
| GBA Bi hGRADE ins2 f | acgacacgtccaactggtggagtctgg | Insert 2 VHH-GFP4 |
| GBA Bi hGRADE ins2 r v2 | aactagaaggcacaggatatcttagctggagacggtgacc | Insert 2 VHH-GFP4 |
| mCherry BamHi f | atatGGATCCTCGCCACCATGGTGAG | mCherry |
| mCherry SpeI r | atatACTAGTCTACTTGTACAGCTCGTCC | mCherry |
| GBA Bi AFB2 ins1 f | tggtaccgtcgacggatccctaatcaagtttgacccgtttggc | Insert 1 AtAFB2-mCherry-weak NLS |
| GBA Bi AFB2 ins1 r | ctcgtgatcgatgaactactttcccgatgagg | Insert 1 AtAFB2-mCherry-weak NLS |
| GBA Bi AFB2 ins2 f | gtagttcatcgatcacgagactagcctcg | Insert 2 pTRE-BI promoter |
| GBA Bi AFB2 ins2 r | gaagcccataattcggggccgcg | Insert 2 pTRE-BI promoter |
| GBA Bi AFB2 ins3 f | cccgaattatgggcttctctgagaccg | Insert 3 miniIAA7 |
| GBA Bi AFB2 ins3 r | caccgatatccttgtcatcgtcatccttgtaatcga | Insert 3 miniIAA7 |
| GBA Bi AFB2 ins4 f | gatgacaaggatatcggtgcaggcgg | Insert 4 VHH-GFP4 |
| GBA Bi AFB2 ins4 r | aggatatctgcagaattcgctagcttagctggagacggtgacct | Insert 4 VHH-GFP4 |
| GBA Bi TIR1 ins1 f | tggtaccgtcgacggatccctaatcaagtttgacccgtttggc | Insert 1 mCherry-weak NLS |
| GBA Bi TIR1 ins1 r | aaaatactgggcggatctggc | Insert 1 mCherry-weak NLS |
| GBA Bi TIR1 ins2 f | gatccgcccagtattttcacgaagtttggagcg | Insert 2 TIR1 |
| GBA Bi TIR1 ins2 r | tcgtgatcgatgacgtacttccctgaagaagt | Insert 2 TIR1 |
| GBA Bi TIR1 ins3 f | tacgtcatcgatcacgagactagcctcg | Insert 3 pTRE-BI promoter |
| GBA Bi TIR1 ins3 r | gtaggccataattcggggccgcg | Insert 3 pTRE-BI promoter |
| GBA Bi TIR1 ins4 f | cccgaattatggcctacccatacgatgt | Insert 4 mAID-VHH-GFP4 |
| GBA Bi TIR1 ins4 r | aggatatctgcagaattcgctagcttagctggagacggtgacct | Insert 4 mAID-VHH-GFP4 |
| GBA Bi TRIM21 ins1 f | tggtaccgtcgacggatccctaatcaagtttgacccgtttggc | Insert 1 mCherry-weak NLS |
| GBA Bi TRIM21 ins1 r | actgactatggcggatctggcgg | Insert 1 mCherry-weak NLS |
| GBA Bi TRIM21 ins2 f | gatccgccatagtcagtggatccttgtgatccaatatt | Insert 2 TRIM21 |
| GBA Bi TRIM21 ins2 r | cgtgatcgATGgcttcagcagcacg | Insert 2 TRIM21 |
| GBA Bi TRIM21 ins3 f | gaagcCATcgatcacgagactagcctcg | Insert 3 pTRE-BI promoter |
| GBA Bi TRIM21 ins3 r | ATCTAACCATaattcggggccgcg | Insert 3 pTRE-BI promoter |
| **Primer** | **Primer Sequence (5'-3')** | **Insert / Amplicon** |

| | | |
|---|---|---|
| GBA Bi TRIM21 ins4 f | ccccgaattATGGTTAGATCTGACAAAACTCACACA | Insert 4 hIgG1 FC2 |
| GBA Bi TRIM21 ins4 r | accgatatcTTTACCCGGAGACAGGGAGAG | Insert 4 hIgG1 FC2 |
| GBA Bi TRIM21 ins5 f | CCGGGTAAAgatatcggtgcaggcgg | Insert 5 VHH-GFP4 |
| GBA Bi TRIM21 ins5 r | aggatatctgcagaattcgctagcttagctggagacgg tgacct | Insert 5 VHH-GFP4 |

Primers for cloning of HDR templates

| Primer | Primer Sequence (5'-3') | Insert / Amplicon |
|---|---|---|
| HDR hSRSF3-GFP Ins1 F | cgcgggaattcgattggacgatgggtgcc | Insert 1 5'-HA |
| HDR hSRSF3-GFP Ins1 R | ggaatatcataatcttttctttcatttgacctag atcgactaaaaaaaaaaaaaaaaaaagaa | Insert 1 5'-HA |
| HDR hSRSF3-GFP Ins2 F | aaatgaaagaaaagattatgatattccaactact gcaagcg | Insert 2 GFP-IRES-NeoR |
| HDR hSRSF3-GFP Ins2 R | caaactgtctttcagaagaactcgtcaagaaggc | Insert 2 GFP-IRES-NeoR |
| HDR hSRSF3-GFP Ins3 F | gttcttctgaaagacagtttgcaagagaagtgg | Insert 3 3'-HA |
| HDR hSRSF3-GFP Ins3 R | cgcgaattcactagtgattgaatcgtgtaaccaa ctgcttcc | Insert 3 3'-HA |
| HDR mSRSF3-GFP Ins1 F | cgcgggaattcgattggtttgcatgggttccaaa tact | Insert 1 5'-HA |
| HDR mSRSF3-GFP Ins1 R | ggaatatcataatcttttctttcatttgacctag atcggct | Insert 1 5'-HA |
| HDR mSRSF3-GFP Ins2 F | aaatgaaagaaaagattatgatattccaactact gcaagcg | Insert 2 GFP-IRES-NeoR |
| HDR mSRSF3-GFP Ins2 R | aactggtctttcagaagaactcgtcaagaaggc | Insert 2 GFP-IRES-NeoR |
| HDR mSRSF3-GFP Ins3 F | gttcttctgaaagaccagtttgcaaaagtgg | Insert 3 3'-HA |
| HDR mSRSF3-GFP Ins3 R | cgcgaattcactagtgattcctgaactggcttca acac | Insert 3 3'-HA |
| HDR hSRSF5-GFP Ins1 F | cgcgggaattcgattggatctcaaagatttcatg agacaagc | Insert 1 5'-HA |
| HDR hSRSF5-GFP Ins1 R | tatcataatcattactgtcaactgatctggacct | Insert 1 5'-HA |
| HDR hSRSF5-GFP Ins2 F | tcagttgacagtaatgattatgatattccaacta ctgcaagcG | Insert 2 GFP-IRES-NeoR |
| HDR hSRSF5-GFP Ins2 R | gttatttacagttcagaagaactcgtcaagaagg c | Insert 2 GFP-IRES-NeoR |
| HDR hSRSF5-GFP Ins3 F | gagttcttctgaactgtaaataacttgccctggg | Insert 3 3'-HA |
| HDR hSRSF5-GFP Ins3 R | cgcgaattcactagtgattggctgaggcaggaga atcg | Insert 3 3'-HA |
| HDR mSRSF5-GFP Ins1 F | cgcgggaattcgattgtgtggtatatgccatgtt tgtactt | Insert 1 5'-HA |
| HDR mSRSF5-GFP Ins1 R | tatcataatcattactgtcaactgatctggacct g | Insert 1 5'-HA |
| HDR mSRSF5-GFP Ins2 F | tcagttgacagtaatgattatgatattccaacta ctgcaagcG | Insert 2 GFP-IRES-NeoR |
| HDR mSRSF5-GFP Ins2 R | gttatttacagttcagaagaactcgtcaagaagg c | Insert 2 GFP-IRES-NeoR |
| HDR mSRSF5-GFP Ins3 F | gagttcttctgaactgtaaataacttgccctggg | Insert 3 3'-HA |
| HDR mSRSF5-GFP Ins3 R | cgcgaattcactagtgattccctgttctacaaat tgagttccaggact | Insert 3 3'-HA |
| **Primer** | **Primer Sequence (5'-3')** | **Insert / Amplicon** |
| HDR hSRRM2-GFP Ins1 F | cgcgggaattcgattcacccagcgccttgc | Insert 1 5'-HA |
| HDR hSRRM2-GFP Ins1 R | atatcataatctggagacctgcaagagaagatat g | Insert 1 5'-HA |

| | | |
|---|---|---|
| HDR hSRRM2-GFP Ins2 F | caggtctccagattatgatattccaactactgca agcG | Insert 2 GFP-IRES-NeoR |
| HDR hSRRM2-GFP Ins2 R | gtggctccatcagaagaactcgtcaagaaggc | Insert 2 GFP-IRES-NeoR |
| HDR hSRRM2-GFP Ins3 F | tcttctgatggagccactgtcccttcttccccag cag | Insert 3 3'-HA |
| HDR hSRRM2-GFP Ins3 R | cgcgaattcactagtgattgggtcccactccctg | Insert 3 3'-HA |
